# Eliminating Mosquitoes with Precision Guided Sterile Males

**DOI:** 10.1101/2021.03.05.434167

**Authors:** Ming Li, Ting Yang, Michelle Bui, Stephanie Gamez, Tyler Wise, Nikolay P. Kandul, Junru Liu, Lenissa Alcantara, Haena Lee, Jyotheeswara R. Edula, Robyn Raban, Yinpeng Zhan, Yijin Wang, Nick DeBeaubien, Jieyan Chen, Hector M. Sanchez C., Jared B. Bennett, Igor Antoshechkin, Craig Montell, John M. Marshall, Omar S. Akbari

## Abstract

The mosquito *Aedes aegypti* is the principal vector for arboviruses including dengue/yellow fever, chikungunya, and zika, infecting hundreds of millions of people annually. Unfortunately, traditional control methodologies are insufficient, so innovative control methods are needed. To complement existing measures, here we develop a molecular genetic control system termed precision guided sterile insect technique (pgSIT) in *Aedes aegypti*. PgSIT uses a simple CRISPR-based approach to generate sterile males that are deployable at any life stage. Supported by mathematical models, we empirically demonstrate that released pgSIT males can compete, suppress, and eliminate mosquitoes in multigenerational population cages. This platform technology could be used in the field, and adapted to many vectors, for controlling wild populations to curtail disease in a safe, confinable, and reversible manner.

## Introduction

Mosquitoes are the world’s deadliest animals, killing more humans than any other animal (*1*) due to transmitting the majority of vector-borne diseases, such as the notorious arboviruses dengue, Zika, yellow fever, and chikungunya transmitted by *Aedes* mosquitoes. The predominating strategy to control these devastating diseases is the use of insecticides, though mosquitoes are evolving and spreading insecticide resistance (*2*), hampering control efforts. Therefore, there is an urgent demand for innovative mosquito-control technologies that are effective, sustainable, and safe.

Alongside traditional control measures, several genetic-based techniques are being used to combat mosquitoes. These include multiple male (♂) release programs aimed at population suppression, such as the classical radiation-based sterile insect technique (SIT), relying on releasing irradiated sterile ♂’s (*3*). Alternative approaches include the *Wolbachia*-based incompatible insect technique (IIT), relying on the release of *Wolbachia* infected ♂’s (*4, 5*), or the antibiotic-based Release of Insects carrying a Dominant Lethal (RIDL) (*6*). Moreover, emerging CRISPR-based homing gene drives that spread target genes through a population faster than through traditional Mendelian inheritance are presently under development with the aim of safe implementation in the future (*7, 8*).

As an alternative, a CRISPR-based technology termed precision-guided SIT (pgSIT), was recently developed in flies (*9*). pgSIT uses a binary approach to simultaneously disrupt genes essential for female (♀) viability and ♂ fertility, resulting in the exclusive survival of sterile ♂’s that can be deployed at any life stage to suppress populations. It requires two breeding strains, one expressing Cas9 and the other expressing guide RNAs (gRNAs). Mating between these strains results in RNA-guided mosaic target gene mutations throughout development, ensuring complete penetrance of desired phenotypes. Compared to alternatives, pgSIT does not require the use of radiation, *Wolbachia*, nor antibiotics, and will not persist in the environment. Unfortunately, this technology is presently only accessible in flies, and for population control techniques, its equivalent needs to be developed for mosquitoes.

To address this, here we systematically engineer pgSIT in *Ae. Aegypti* using a system that simultaneously disrupts genes essential for ♂ fertility and ♀flight, which is necessary for mating, blood feeding, reproduction, and predator avoidance—meaning survival in general (*10*). Using our technology, we demonstrate resulting progeny of flightless ♀’s and fit sterile ♂’s that can compete, suppress, and eliminate mosquito populations in multigenerational population cages. Mathematical models suggest that releases of *Ae. aegypti* pgSIT eggs could effectively eliminate a local *Ae. aegypti* population using achievable release schemes. Taken together, this study suggests pgSIT may be an efficient technology for mosquito population control and the first example of one suited for real-world release.

## Results

### Validation of pgSIT target genes

To engineer pgSIT in *Ae. aegypti*, we first validated target genes by generating transgenic gRNA-expressing lines targeting two conserved genes: *β-Tubulin 85D* (*βTub*, AAEL019894), specifically expressed in mosquito testes (*11*–*13*) and essential for spermatogenesis and ♂ fertility (*14*), and *myosin heavy chain* (*myo-fem*, AAEL005656), expressed nearly exclusively in ♀pupae (*11, 12*) and essential for ♀flight (*15*) (**Fig. S1, Table S1**). To ensure efficient disruption, each gRNA line encoded four U6–promoter-driven (*16*) gRNAs targeting unique sites in the coding sequence of either *βTub (U6-gRNA*^*βTub*^ *-* marked with 3xP3-GFP) or *myo-fem (U6-gRNA*^*myo-fem*^ - marked with 3xP3-tdTomato) (**Fig. S2-4**). Multiple independent transgenic lines were generated, and to assess their activity, we conducted bidirectional crosses with Cas9 controlled by a homozygous *nuclear pore complex protein* (*Cas9 -* marked with Opie2-CFP) (*17*) (**Table S2, S3, Fig. S2-4)**. The resulting transheterozygous F_1_ progeny (*gRNA*/+, Cas9/+) were assessed and crossed to wildtype (WT) for further evaluation. For the *βTub* crosses, fertility of the F_1_ transheterozygous ♂’s ranged from 0–94.9%, with two lines achieving 100% sterility from immotile sperm (*14*), while F1 transheterozygous ♀’s maintained normal fertility (**Fig. 1, S2, Table S3, Video S1**). For *myo-fem* crosses, all F1 transheterozygous ♀’s generated from ⅗ lines were flightless, while F1 transheterozygous ♂’s maintained normal flight (**Fig. 1, Table S3, Fig. S3, Video S2**,**3**). As expected, ♀flightlessness significantly reduced mating ability and blood consumption as many get trapped on the water surface following eclosion, resulting in reduced fecundity, fertility, and survival. Sanger sequencing of genomic DNA revealed expected mutations at the *βTub-* and *myo-fem*-targeted loci.

**Figure 1.**
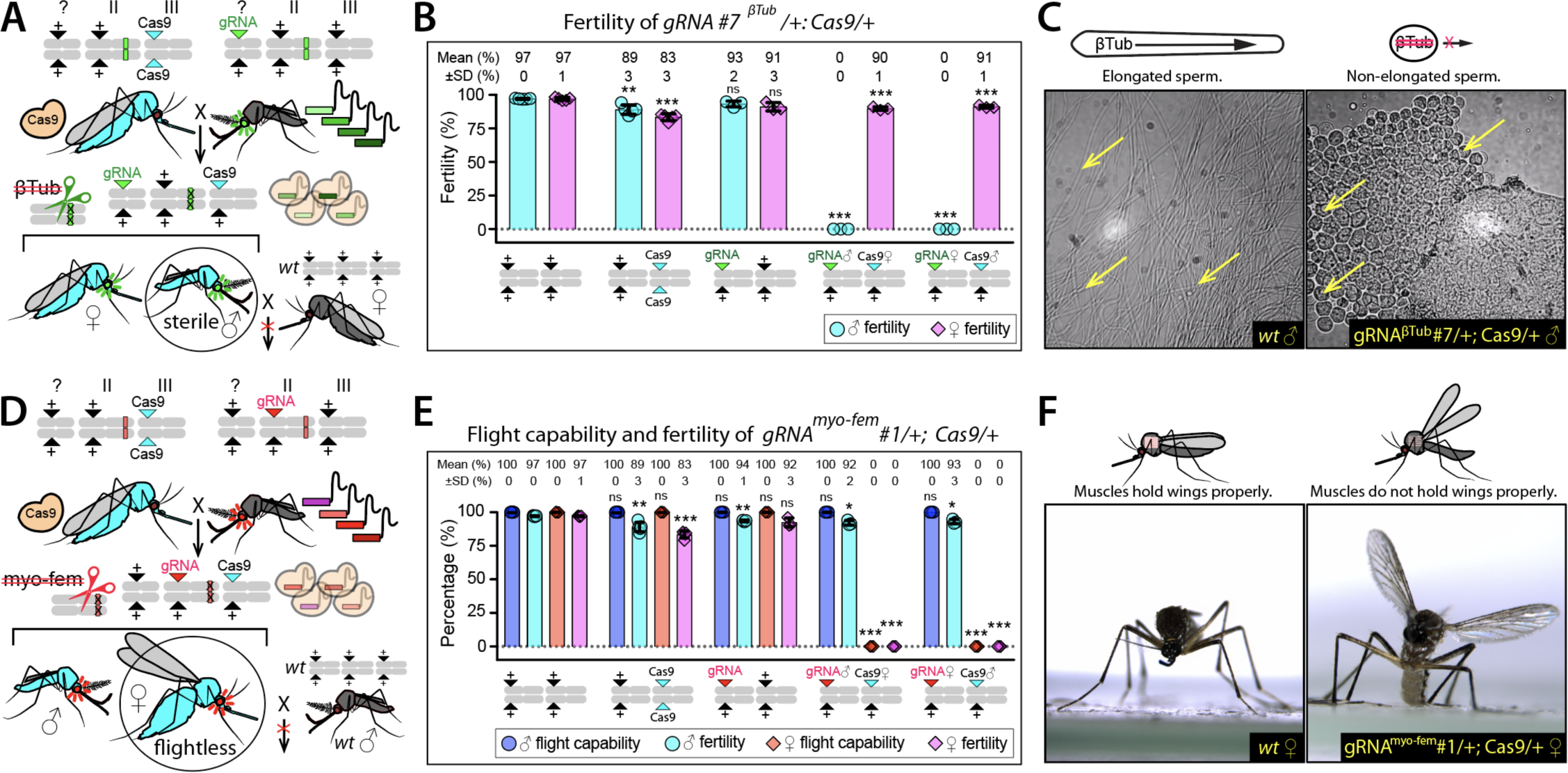
Validation of pgSIT target genes. Cas9/gRNA-mediated disruption of *βTub* or *myo-fem* results in (**A**-**C**) male (♂) sterility or (**D**-**E**) female (♀) flightlessness, respectively. Schematics of genetic crosses to assess the efficiency of (**A**) *βTub* or (**D**) *myo-fem* disruption in the F_1_ transheterozygous progeny. (**B**) Histogram indicating the percent of fertile progeny for each of the various progeny genotypes using *gRNA*^*βTub*^#7 line. **(C)** Imaging of seminal fluid from WT and *gRNA*^*βTub*^*#7/+;Cas9/+* mosquitoes, showing the difference in spermatid elongation caused by the disruption of *βTub* (**Video S1**). **(E)** Histogram showing percent of fertile and flight-capable mosquitoes in each cross using *gRNA*^*myo-fem*^*#1* line. (**Video S1-S3). (F)** Imaging showing the specific wing posture phenotype induced by the *myo-fem* disruption in females, but not in males, in which the resting wings were uplifted. Data from both paternal Cas9 crosses (Cas9♂ ×gRNA♀) and maternal Cas9 crosses (Cas9♀×gRNA♂) are shown (**Fig. S2, S3, Table S3**). Bar plots show means ± one standard deviation (SD), biological replicates, and mean and SD values rounded to a whole number. Statistical significance was estimated using a two-sided Student’s *t* test with unequal variance. (*p* ≥ 0.05^ns^, *p* < 0.05*, *p* < 0.01**, and *p* < 0.001***).

### Development of pgSIT and fitness assessments

To generate a pgSIT strain capable of targeting both *βTub* and *myo-fem* simultaneously, we combined two gRNA lines that exclusively produced sterile ♂’s (*gRNA*^*βTub*^*#7*) or flightless ♀’s (*gRNA*^*myo-fem*^*#1*) (**Fig. 1, Fig. S2-S3, Table S3**) by repeated introgression, generating a trans-homozygous stock (termed *gRNA*^*βTub*+*myo-fem*^) (**Fig. S4**). To assess its activity, we bidirectionally crossed *gRNA*^*βTub*+*myo-fem*^ to *Cas9*. Importantly, these crosses yielded all flightless ♀’s (termed *pgSIT*^*♀*^) and sterile ♂’s (termed *pgSIT*^*♂*^) with normal flight and mating capacity (**Fig. 2, S5, Table S4-8, Video S4-6**). We next determined transgene integration sites, single copy number per transgene, and confirmed target gene disruptions by both amplicon sequencing (**Fig. S6**) and Nanopore genome sequencing using transheterozygous *pgSIT*^*♂’*^s (**Fig. S7-S9, Table S9-10**). We also performed transcriptome sequencing of pupae comparing *pgSIT*^*♂’*^s and *pgSIT*^*♀’*^s to WT to quantify target gene reduction, expression from transgenes, and to assess global expression patterns (**Fig. S8-S10, Table S11-S15**). As expected, we observed significant target gene disruption in *pgSIT* individuals, robust expression from our transgenes, and non-target gene misexpression, which would be expected given the significant phenotypes observed (i.e. flightless females and spermless males).

**Figure 2.**
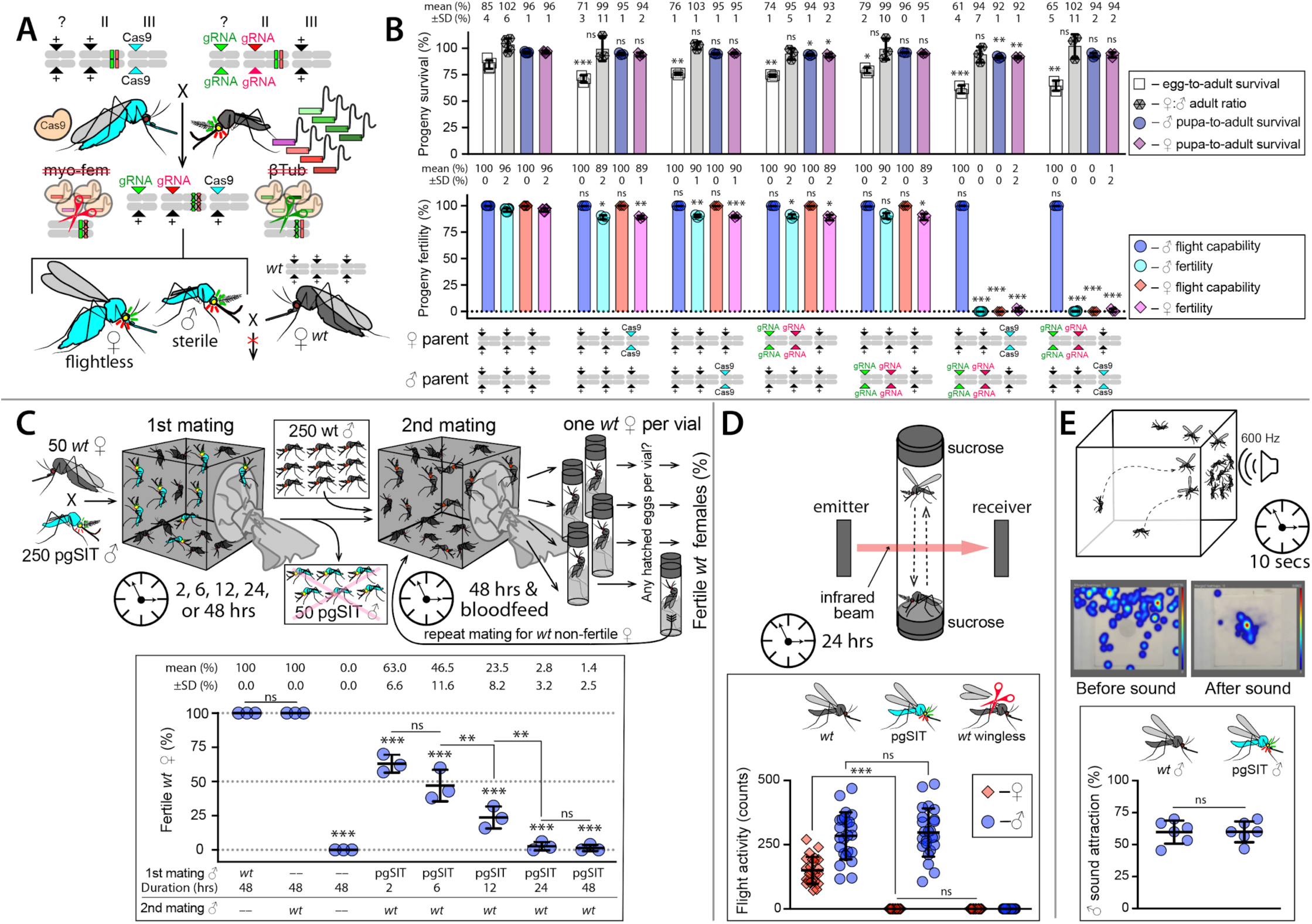
Genetic characterization of pgSIT. (**A**) The pgSIT cross between transhomozygous gRNA ♂’s harboring both *gRNA*^*βTub*^*#7* and *gRNA*^*myo-fem*^*#1* (termed: *gRNA*^*βTub*+*myo-fem*^) and the homozygous *Cas9*. The pgSIT cross was initiated reciprocally to generate F_1_ transheterozygous progeny carrying either maternal or paternal Cas9. (**B**) Histogram comparing the survival and fitness of transheterozygous and heterozygous Cas9 or gRNA progeny to those of WT (**Table S4, Video S4**).(**C**) Experimental set-up to determine whether prior matings with *pgSIT*^*♂*^’s suppresses WT ♀fertility. WT ♀’s were cohabitated with *pgSIT*^*♂*^’s for 2, 6, 12, 24, or 48 hours then WT ♀’s were transferred to a new cage along with WT **♂’**s and mated for an additional 2 days. The ♀’s were then blood fed and individually transferred to a vial. Eggs were collected and hatched for fertility determination. Following this, non-fertile ♀’s were then placed back into cages along WT ♂’s for another chance to produce progeny. This was repeated for up to five gonotrophic cycles, and the percentage of fertile ♀’s in each group of 50 ♀’s was plotted (**Table S8**). (**D**) Flight activity of individual mosquitoes including was assessed for 24 hours using a vertical *Drosophila* Activity Monitoring (DAM) System, which uses an infrared beam to record flight (**Table S6, Video S5**). (**E**) To quantify the attractiveness of ♂’s to ♀’s for mating, we used a mating-behavior lure of a tone mimicking ♀flight. A 10-second 600 Hz sine tone was applied on one side of the cage, and a number of mosquito **♂’**s landing on the mesh around a speaker was scored. Heatmaps were generated using Noldus Ethovision XT. (**Table S7, Video S6**). Plots show biological replicates and means ± SDs. Statistical significance was estimated using a two-sided Student’s *t* test with unequal variance. (*p* ≥ 0.05^ns^, *p* < 0.05*, *p* < 0.01**, and *p* < 0.001***).

To explore potential fitness effects, we assayed several fitness parameters including ♀fecundity, fertility, flight activity, ♂ mating capacity, ♂ sound attraction, larva-pupa development time, pupa-adult development time, and longevity (**Fig. 2, S5, Table S5-8, Video S5-6**). The *pgSIT*^*♀*^’s were flightless with significantly reduced fecundity, fertility, and survival, indicating they would be very unlikely to survive in the wild, let alone transmit pathogens. For *pgSIT*^*♂*^’s, other than slightly delayed larva-pupa development time, we did not detect significant differences in fitness parameters. Previous studies demonstrated that *Ae. aegypti* ♀’s typically mate only once in their lifetime, a behavior known as monandry (*18*). To explore whether prior matings with *pgSIT*^*♂*^’s could suppress ♀fertility, we initiated experiments in which WT ♀’s were first mated with *pgSIT*^*♂*^’s for a period of time (2, 6, 12, 24, or 48 hrs) followed by WT ♂’s (48 hrs). Fertility was measured for up to five gonotrophic cycles. We found that prior exposure to *pgSIT*^*♂*^’s ensured long lasting reductions in ♀fertility, spanning 5 gonotrophic cycles, with longer exposures (24 and 48 hrs) resulting in near complete suppression of ♀fertility (**Fig. 2, Table S8**).

### pgSIT induced population suppression

To assess whether *pgSIT* ♂’s could compete and suppress populations, we conducted discrete, multi-generational, population cage experiments by repeatedly releasing either eggs or adult ♂’s each generation, using several introduction frequencies (pgSIT:WT - 1:1, 5:1, 10:1, 20:1, and 40:1) (**Fig. 3, Table S16**). To measure efficacy each generation, we counted the total number of eggs laid and hatched and confirmed the lack of presence of marker genes in hatched larvae indicating released *pgSIT* ♂’s were indeed sterile. Adult releases at high release thresholds (20:1, 40:1) eliminated all populations by generation 3, and at lower release thresholds (10:1), we saw elimination by generation 6. For the egg releases, elimination was achieved by generation 6 for 4/6 populations at high release thresholds (20:1, 40:1).

**Figure 3.**
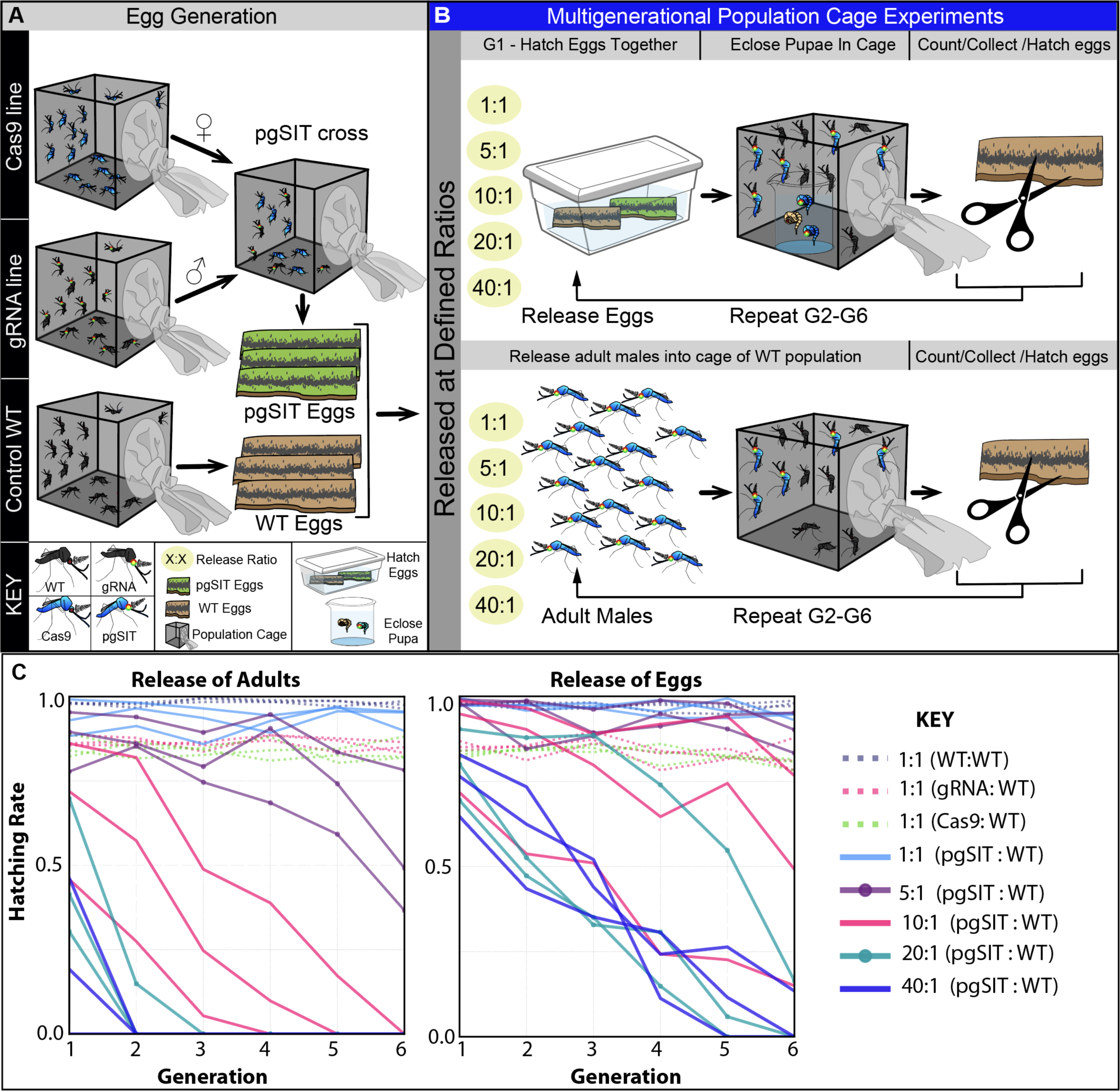
Multigenerational cage trials demonstrating efficient population suppression. **(A)** To generate sufficient mosquitoes, three lines were raised separately, including homozygous Cas9, transhomozygous *gRNA*^*βTub*+*myo-fem*^, and WT. To generate pgSIT progeny, virgin Cas9 ♀’s were genetically crossed to *gRNA*^*βTub*+*myo-fem*^ ♂’s, and eggs were collected. **(B)** To perform multigenerational population cage trials of pgSIT, two strategies were employed: release of eggs (**B**, top panel); release mature adults (**B**, bottom panel). For both strategies, multiple pgSIT:WT release ratios were tested, including: 1:1, 5:1, 10:1, 20:1, and 40:1. Each generation, total eggs were counted, and 100 eggs were selected randomly to seed the subsequent generation. The remaining eggs were hatched to measure hatching rates and score transgene markers. This procedure was repeated after each generation until each population was eliminated (**Table S7)**. **(C)** Multigeneration population cage data for each release threshold plotting the proportion of eggs hatched each generation.

### Theoretical performance of pgSIT in a wild population

To explore the potential for *pgSIT* ♂’s to suppress *Ae. aegypti* populations in the wild, we simulated releases of *pgSIT* eggs on the island of Onetahi, Tetiaroa, French Polynesia (**Fig. 4**), a field site for releases of *Wolbachia*-infected ♂ mosquitoes, using the MGDrivE simulation framework (*19*). Weekly releases of up to 400 *pgSIT* eggs per wild adult were simulated in each human structure over 10-24 weeks. The scale of these releases was chosen considering adult release ratios of 10:1 are common for sterile male mosquito interventions (*6*) and female *Ae. aegypti* produce >30 eggs per day in temperate climates (*20*). We also assumed 25% reductions in male mating competitiveness and adult lifespan for *pgSIT* males by default because, although *pgSIT* fitness effects were not apparent from laboratory experiments, they may become apparent in the field. Results from these simulations suggest that significant population suppression (>96%) is seen for a wide range of achievable release schemes, including 13 weekly releases of 120 or more *pgSIT* eggs per wild adult (**Fig. 4, Video S7**). Population elimination was common for larger yet achievable release schemes, including 18 weekly releases of 200 or more *pgSIT* eggs per wild adult, and 24 weekly releases of 100 or more *pgSIT* eggs per wild adult. Results also suggest a wider range of *pgSIT* fitness profiles (e.g. a 50% reduction in male mating competitiveness and 25% adult lifespan reduction) could lead to population elimination for these release schemes (**Fig. 4**).

**Figure 4.**
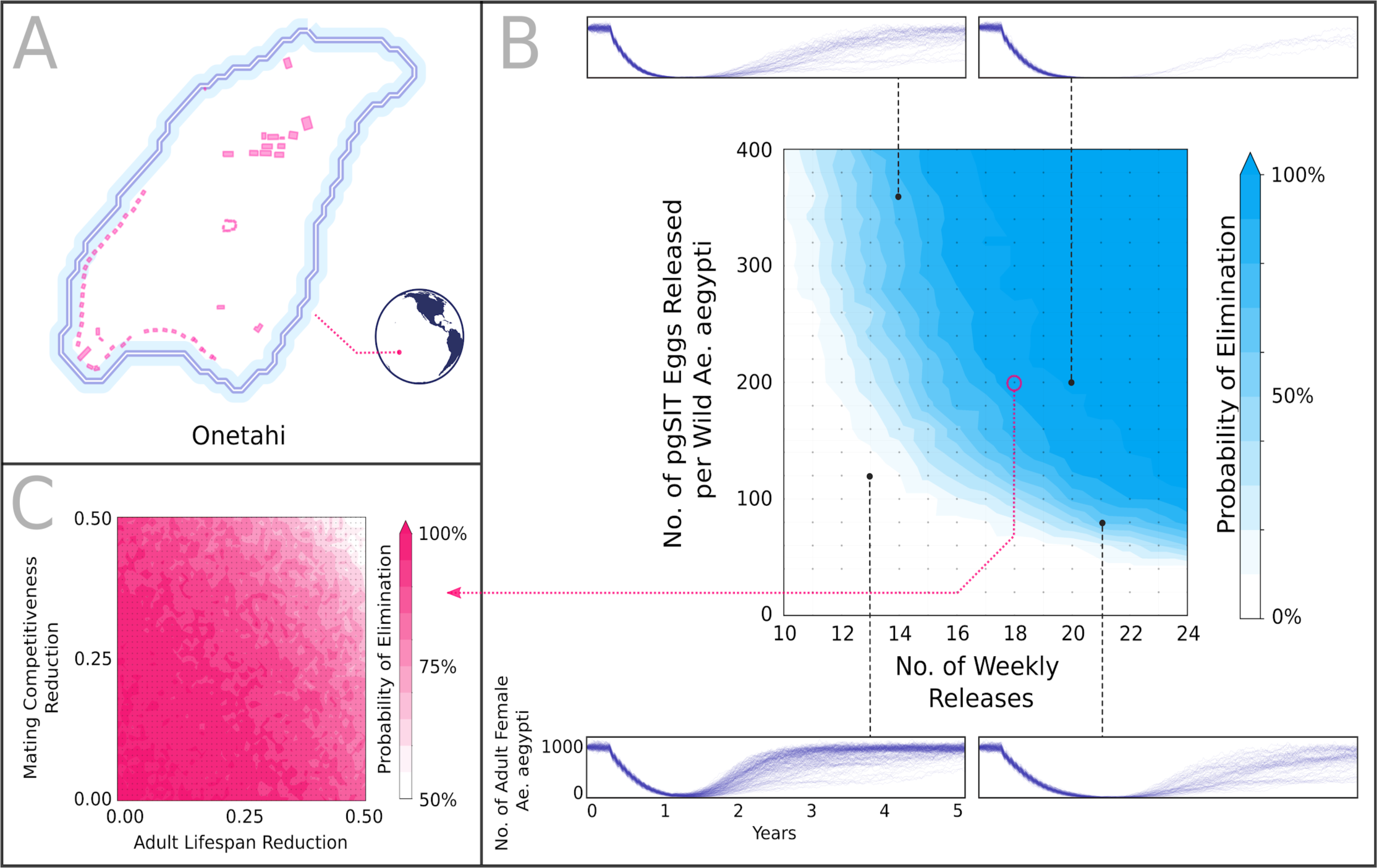
Model-predicted impact of releases of *pgSIT* eggs on *Ae. aegypti* population density and elimination. **(A)** Releases were simulated on the island of Onetahi, Tetiaroa, French Polynesia, a field site for releases of *Wolbachia*-infected ♂ mosquitoes, using the MGDrivE simulation framework (*19*) and parameters described in **Table S17**. Human structures are depicted and were modeled as having an equilibrium population of 16 adult *Ae. aegypti* each. **(B)** Weekly releases of up to 400 *pgSIT* eggs per wild *Ae. aegypti* were simulated in each human structure over 10-24 weeks. The *pgSIT* construct was conservatively assumed to decrease male mating competitiveness by 25% and adult lifespan by 25%. Elimination probability was calculated as the percentage of 200 stochastic simulations that resulted in local *Ae. aegypti* elimination for each parameter set. Sample time-series depicting female *Ae. aegypti* population density are depicted above and below the heatmap. **(C)** Elimination probability (given 18 weekly releases of 200 *pgSIT* eggs per wild *Ae. aegypti*) is depicted for a range of *pgSIT* ♂ fitness profiles. Elimination is possible for a wide range of reductions in male mating competitiveness (0-50%) and adult lifespan (0-50%) for an achievable release scheme.

## Discussion

While many technologies for halting the spread of deadly mosquito-borne pathogens exist, none are without significant drawbacks such that additional measures are needed. By disrupting essential genes throughout development, we demonstrate efficient production of short-lived, flightless *pgSIT*^*♀*^’s and fit sterile *pgSIT*^*♂*^’s. Importantly, when repeatedly released into caged populations, the *pgSIT*^*♂*^’s competed with WT ♂’s thereby suppressing, and even eliminating, populations using release ratios that are achievable in the field (*4*–*6*). Mathematical models suggest that population elimination could be accomplished in the field through sustained releases of ∼100-200 or more *pgSIT* eggs per wild *Ae. aegypti* adult, even if fitness costs significantly exceed those measured in contained laboratory experiments.

For pgSIT to be realized in the wild, the two strains will first need to be separately and continuously mass-reared in a facility, without contamination, and crossed to produce sterile ♂’s. While this can be viewed as rate-limiting (*21*), it offers stability, as the binary CRISPR system will remain inactive until crossed—thereby reducing the evolution of suppressors or mutations that could disrupt the system. Additionally, each sorted ♀can produce up to 450 eggs in her lifetime (*22*), which improves scalability. Moreover, once crossed, the resulting progeny are essentially dead-ends (i.e. sterile ♂’s /flightless ♀’s), hatched among high numbers of sterile *pgSIT*^*♂’*^s, and should not contribute to the gene pool (*23*). We demonstrate here that the technology is fully penetrant by screening >100K individuals.

pgSIT offers an alternative approach to scalability that should help decrease costs and increase efficiency. For instance, the required genetic cross at scale can be initiated using existing robotic sex sorting devices (www.senecio-robotics.com) or (*5*). Upon sex sorting and crossing, the resulting *pgSIT* progeny can be distributed and released at any life stage, mitigating requirements for sex separation at field sites. This strategy will be especially effective for mosquitoes that diapause during the egg stage (e.g. *Aedes* species) because it will enable long-term egg accumulation. Eggs could be distributed to logistically spaced remote field sites where they can hatch, develop, and compete with wild mosquitoes (**Fig. S11**). This attractive feature should reduce the costs of developing multiple production facilities requiring on-site sex separation for manual release of fragile adults.

It should be noted that the releases of adult *pgSIT*^*♂’*^s unexpectedly resulted in faster population suppression as compared to egg releases in multigenerational population cage experiments. We believe this to result from the slightly reduced egg hatching rates of *pgSIT*^*♂’*^s and their delayed larva-pupa development time, which likely enabled the co-released WT ♂’s first access to WT ♀’s. While this could impact the discrete generation population cage experiments conducted here, it should not be problematic for suppressing continuous populations in the wild.

Finally, notwithstanding its inherently safe nature, pgSIT requires genetic modification, and regulatory use authorizations will need to be granted prior to implementation. While this could be viewed as a limitation (*21*), we don’t expect obtaining such authorizations to be insurmountable. In fact, we envision pgSIT to be regulated in a similar manner to Oxitec’s RIDL technology, which has been successfully deployed in many locations and recently received experimental use authorizations in the USA.

Overall, the inherent self-limiting nature of pgSIT, offers a controllable safe alternative to technologies that can persist and spread in the environment, such as gene drives (*8*). Going forward, pgSIT may provide an efficient, safe, scalable, and environmentally friendly alternative next-generation technology for wild population control of mosquitoes resulting in wide-scale prevention of human disease transmission.

## Supplementary materials

### Mosquito rearing and maintenance

*Ae. aegypti* mosquitoes were derived from the Liverpool strain (wildtype [WT]) previously used to generate the reference genome (*24*). Mosquitoes were raised in incubators at 27.0°C with 20–40% humidity and a 12-hour light/dark cycle in cages (Bugdorm, 24.5 cm × 24.5 cm × 24.5 cm). Adults were provided 0.3 M aqueous sucrose *ad libitum*, and ♀’s were blood fed on anesthetized mice for two consecutive days for ∼15 minutes at a time. Oviposition substrates were provided ∼3 days following the second blood meal. Eggs were collected and aged for ∼4 days to allow for embryonic development, then were hatched in deionized H_2_O in a vacuum chamber. Roughly ∼400 larvae were reared in plastic containers (Sterilite, 34.6 cm × 21 cm × 12.4 cm, USA) with ∼3 liters of deionized H_2_O, and fed fish food (TetraMin Tropical Flakes, Tetra Werke, Melle, Germany). For genetic crosses, to ensure ♀virginity, pupae were separated and sexed under the microscope by sex-specific morphological differences in the genital lobe shape (at the end of the pupal abdominal segments just below the paddles) before being released to eclose in cages. These general rearing procedures were followed unless otherwise noted. Mosquitoes were examined, scored, and imaged using the Leica M165FC fluorescent stereo microscope equipped with the Leica DMC2900 camera. For higher resolution images, we used a Leica DM4B upright microscope equipped with a VIEW4K camera enabling time lapse videos. Time lapse videos of caged adult mosquitoes were taken with a mounted Canon EOS 5D Mark IV using a 24–105mm image stabilizer ultrasonic lens.

### Guide RNA design and testing

Two target genes were selected for gRNA design: *β-Tubulin 85D* (*βTub*, AAEL019894) and *myosin heavy chain* (*myo-fem*, AAEL005656). For each target gene, DNA sequences were first identified using reference genome assembly (*24*), and genomic target sites were validated using PCR amplification and Sanger sequencing (**Table S18** for primer sequences). Gene structures, transcripts, and exon-intron junction boundaries were carefully evaluated using comprehensive developmental transcriptome data (*11, 24*) loaded into an internal genome browser. Target gRNA sequences were selected to be 20 bp (N20) in length, excluding the PAM (NGG) (*25*). For *in silico* gRNA selection, we used either CHOPCHOP V3.0.0 (https://chopchop.cbu.uib.no) or CRISPOR (http://crispor.tefor.net) to minimize potential genomic off-target cleavage events. In total, we designed four gRNAs targeting *βTub* and four gRNAs targeting *myo-fem* (**Table S18**). To confirm gRNA activity *in vivo*, each gRNA was *in vitro* synthesized prior to construct design (Synthego, CA, USA). Then 100 ng/ul of gRNA was individually injected into fifty preblastoderm stage embryos (0.5–1 hr old) derived from Exu-Cas9 maternally depositing mothers, per previous embryo-injection protocols (*16, 17*). The surviving G0 progeny were pooled (2– 5 individuals per pool), and genomic DNA was extracted using the DNeasy blood and tissue kit (Qiagen, Cat No./ID: 69506) following the manufacturer’s protocols. To molecularly characterize the induced mutations, target loci were PCR amplified from extracted genomic DNA, and the PCR products were gel purified (Zymo Research, Zymoclean Gel DNA Recovery Kit, Cat No./ID: D4007). The purified products were either sent directly for sequencing or subcloned (Invitrogen, TOPO-TA, Cat No./ID: LS450641), wherein single colonies were selected and cultured in Laurel Broth (LB) with ampicillin before plasmid extraction (Zymo Research, Zyppy plasmid miniprep kit, Cat No./ID: D4036) and Sanger sequencing. Mutated alleles were identified *in silico* by alignment with WT target sequences. All primers used for PCR and sequencing, including gRNA target sequences, are listed in **Table S18**.

### Construct molecular design and assembly

The Gibson enzymatic assembly method was used to engineer all constructs in this study (*26*). To generate the Nup50-Cas9 construct marked with CFP, OA-874PA (Addgene #164846), we used our previous plasmid for Cas9 expression (Addgene plasmid #100608) as the backbone (*17*). The fragments of T2A-eGFP-P10-3’UTR and OpIE2-dsRed-SV40 were removed by cutting with restriction enzyme FseI. Then, the P10-3’UTR fragment was amplified from Addgene plasmid #100608 with primers 874-P10 and 777B. Another fragment, OpIE2-CFP-SV40, was synthesized using gBlocks® Gene Fragment service (Integrated DNA Technologies, Coralville, Iowa). Both fragments were provided for the Gibson assembly into the cut backbone. We designed two constructs, OA-1067A1 (Addgene #164847) and OA-1067K (Addgene #164848), each carrying four different gRNAs targeting either β-Tubulin 85D (*βTub*, AAEL019894) or myosin heavy chain (*myo-fem*, AAEL005656) genes.

To engineer these plasmids, four intermediate plasmids, OA-1055A (*gRNA*^*βTub1&2*^), OA-1055B (*gRNA*^*βTub3&4*^), OA-1055W (*gRNA*^*myo-fem1&2*^), and OA-1055X (*gRNA*^*myo-fem3&4*^), each harboring two gRNAs, were generated by cutting a backbone plasmid OA-984 (Addgene plasmid #120363), which contains piggyBac elements and the 3xP3-tdTomato transformation marker, with the restriction enzymes AvrII and AscI. Two gBlocks® Gene Fragments were then cloned in, each containing two gRNAs: one driven by U6b (AAEL017774) and one by U6c (AAEL017763) promoters (*17*). To assemble the final plasmid OA-1067A1, an intermediate plasmid OA-1067A was generated by linearizing the plasmid OA-1055B with the restriction enzyme BglII and inserting in the fragment of U6b-*gRNA*^*βTub1*^-U6c-*gRNA*^*βTub2*^ amplified with primers 1167.C1 and 1067.C2 from plasmid OA-1055A. Then, the fragment of 3xP3-tdTomato was removed from plasmid OA-1067A using the restriction enzymes AscI and NotI and replaced with the 3xP3-eGFP transformation marker amplified with primers 1067A1.C1 and 1067A1.C2 from the plasmid OA-961B (Addgene plasmid #104967). To assemble the final plasmid OA-1067K, OA-1055W was linearized with the restriction enzyme FseI, and the insertion of U6b-*gRNA*^*myo-fem3*^-U6c-*gRNA*^*myo-fem4*^ was amplified with primers 1167.C5 and 1067.C6 from the plasmid OA-1055X. During each cloning step, single colonies were selected and cultured in LB medium with ampicillin, and then the plasmids were extracted (Zymo Research, Zyppy plasmid miniprep kit, Cat No./ID: D4036) and Sanger sequenced. Final plasmids were maxi-prepped using (Zymo Research, ZymoPURE II Plasmid Maxiprep kit, Cat No./ID: D4202) and Sanger sequenced. All primers are listed in **Table S18**. Complete plasmid sequences and plasmid DNA are available at www.addgene.com.

### Generation of transgenic lines

Transgenic lines were generated by microinjecting preblastoderm stage embryos (0.5–1 hr old) with a mixture of the piggybac plasmid (200 ng/ul) and a transposase helper plasmid (phsp-Pbac, (200 ng/ul). Embryonic collection and microinjections were performed following previously established procedures (*17*). After 4 days of development post-microinjection, G0 embryos were hatched in deionized H_2_O in a vacuum chamber. Surviving G0 pupae were separated and sexed and divided into separate ♀or ♂ cages (∼20 cages total). The pupae eclosed inside these cages along with added WT ♂ pupae (added into the ♀cages) or WT ♀pupae (added into the ♂ cages) at 5:1 ratios (WT:G0). Several days post-eclosion (∼4–7), enabling sufficient time for development and mating, a blood meal was provided, and eggs were collected, aged, then hatched. The hatched larvae with positive fluorescent markers were individually isolated using a fluorescent stereo microscope (Leica M165FC). To isolate separate insertion events, selected transformants were individually crossed to WT (5:1 ratios of WT:G1), and separate lines were established (**Table S2**). These were subjected to many generations of backcrosses to WT to isolate single insertion events. Each of these individual gRNA lines (OA-1067A1: *gRNA*^*βTub*^ and OA-1067K: *gRNA*^*myo-fem*^) were maintained as mixtures of homozygotes and heterozygotes with periodic selective elimination of WTs. The Cas9 line (OA-874PA: Nup50-Cas9) was homozygosed by ∼10 generations of single-pair sibling matings selecting individuals with the brightest expressing transformation markers. Homozygosity was confirmed genetically by repeated test crosses to WT.

### Genetic testing of established lines

To assess the activity of the transgenic lines generated, we performed a series of genetic crosses by releasing sexed pupae into cages. We first crossed gRNA lines (*gRNA* ♂ ×WT ♀) to generate heterozygotes. We next reciprocally crossed heterozygous *gRNA*^*βTub*^/+ (lines #1–10) and the heterozygous *gRNA*^*myo-fem*^/+ (lines #1–5), with homozygous Cas9 (1 ♂ ×10 ♀). To measure the fecundity, the resulting transheterozygous F_1_ progeny (*gRNA*^*βTub*^/+; Cas9/+), or (*gRNA*^*myo-fem*^/+; Cas9/+), were reciprocally crossed to WT’s (50 ♂ ×50 ♀), keeping track of the grandparents genotypes (**Fig. S2, S3, Table S3**). Control crosses of: WT ♂ ×WT ♀; WT ♂ ×Cas9 ♀; Cas9 ♂ ×WT ♀; *gRNA*/+ ♂ ×Cas9 ♀; *gRNA*/+ ♀×Cas9 ♂; *gRNA*/+ ♀×WT ♂; and *gRNA*/+ ♂ ×WT ♀were also set up for comparisons (50 ♂ ×50 ♀). Adults were allowed to mate in the cage for 4–5 days, then blood meals were provided, and eggs were collected and hatched. The percentage of egg hatching (i.e. fertility) was estimated by dividing the total number of eggs laid by the total number of hatched eggs. Larvae-to-adult survival rates were calculated by dividing the total number of adults that emerged by the total number of larvae. Pupae-adult survival rates were calculated by dividing the number of dead pupae by the total number of pupae. Flight capacity for each sex was calculated by dividing the total number that were flightless (observed by eye) by the total of number of adult mosquitoes of that sex. Blood acquisition rates were calculated by dividing the number of blood-fed ♀’s by the total number of ♀’s. To investigate ♂ internal anatomical features, testes and ♂ accessory glands (n = 20) were dissected in 1% PBS buffer for imaging.

### Generation and characterization of *gRNA*^*βTub*+*myo-fem*^

To generate *gRNA*^*βTub*+*myo-fem*^, we genetically crossed *gRNA*^*βTub#7*^ (marked with 3xp3-GFP) with *gRNA*^*myo-fem#1*^(marked with 3xp3-tdTomato). Resulting F1 transheterozygotes *gRNA*^*βTub#7*^ /+; *gRNA*^*myo-fem#1*^ /+ were subjected to multiple generations of single-pair sibling matings, carefully selecting individuals with the brightest expressing transformation markers, to generate a transhomozygous stock (termed: *gRNA*^*βTub*+*myo-fem*^). Zygosity was confirmed genetically by repeated test crosses to WT. To measure efficacy, we bidirectionally crossed *gRNA*^*βTub*+*myo-fem*^ with Cas9 (50 ♂ ×50 ♀), generating F1 transheterozygotes *gRNA*^*βTub*+*myo-fem*^/+; Cas9/+. Control crosses were also setup for comparisons: *gRNA*^*βTub*+*myo-fem*^ ♂ ×*gRNA*^*βTub*+*myo-fem*^ ♀; *gRNA*^*βTub*+*myo-fem*^ ♂ ×WT ♀; *gRNA*^*βTub*+*myo-fem*^ ♀×WT ♂; Cas9 ♂ ×Cas9 ♀; Cas9 ♂ ×WT ♀; and Cas9 ♀×WT ♂; (50 ♂ ×50 ♀). To determine the fecundity and fertility, resulting transheterozygous F1’s (∼3 days old) were bidirectionally crossed to WT’s (50 ♂ ×50 ♀; 10 replicates each). These were allowed to mate for ∼2 days and then blood fed. Afterwards, eggs were collected for up to five consecutive gonotrophic cycles and hatched.

### Determination of transgene integration sites and copy number

To determine the transgene insertion site(s) and copy number(s), we performed Oxford Nanopore DNA sequencing. We extracted genomic DNA using the Blood & Cell Culture DNA Midi Kit (Qiagen, Cat# 13343) from twenty adult transheterozygous *pgSIT*^*♂’*^s (3 days old) harboring all three transgenes (Cas9/+; *gRNA*^*βTub#7*^ /+; *gRNA*^*myo-fem#1*^ /+), following the manufacturer’s protocol. The sequencing library was prepared using the Oxford Nanopore SQK-LSK109 genomic library kit and sequenced on a single MinION flowcell (R9.4.1) for 72 hrs to generate an N50 read length for the set of 4088 bp. Basecalling was performed using ONT Guppy basecalling software version 4.4.1, generating 2.94 million reads above quality threshold Q≧7, which corresponds to 8.68 Gb of sequence data. To determine transgene copy number(s), reads were mapped to the AaegL5.0 reference genome (*24*) supplemented with transgene sequences (OA-1067A1: *gRNA*^*βTub*^; OA-1067K: *gRNA*^*myo-fem*^; and OA-874PA: Nup50-Cas9) using minimap2 (*27*). In total, 2,862,171 out of 2,936,275 reads (97.48%) were successfully mapped with a global genome-wide depth of coverage of 5.495. We calculated the mean coverage depth for all contigs in the genome (2310) and the three plasmids (OA-1067A1: *gRNA*^*βTub*^; OA-1067K: *gRNA*^*myo-fem*^; and OA-874PA: Nup50-Cas9) as well as normalized coverage (**Table S9-S10**). Transgene coverage ranged from 5.1 to 7.6, and normalized coverage ranged from 0.93 to 1.38. As compared to the three chromosomes, the coverages are consistent with the transgenes present at a single copy (**Fig. S7**).

To identify transgene insertion sites, we inspected reads that aligned to the transgenes in the Interactive Genomics Viewer (IGV) browser. The reads extending beyond the boundaries of the transgenes were then analyzed to determine mapping sites within the genome. For OA-874PA, one read spanned the whole transgene (∼11.5 kb) and extended 4 and 3.5 kb on both sides. The extending portions mapped to both sides of the position on NC_035109.1:33,210,105 (chromosome 3), with the nearest gene being AAEL023567, which is ∼5 kb away. For OA-1067K, one read covered ∼7 kb of the transgene extending ∼10 kb off the 3’ end, 9 kb of which map to the NC_035108.1:287,686-296,810 region (chromosome 2). A few other shorter reads map to the same location. The site is located in the intron of AAEL005206, which is a capon-like protein, and based on the RNA-seq data, it’s expression does not appear to be affected in pgSIT animals. For OA-1067A1, the nanopore sequencing was unable to resolve the insertion site, presumably due to its insertion in one of the remaining gaps in the genome. Finally, using nanopore data, we confirmed genomic deletions in both pgSIT target genes - see AEL019894 and AAEL005656 as expected (**Fig. S8-S9**). The nanopore sequencing data has been deposited to the NCBI sequence read archive (SRA) under BioProject ID is PRJNA699282 with accession number SRR13622000.

### Transcriptional profiling and expression analysis

To quantify target gene reduction and expression from transgenes as well as to assess global expression patterns, we performed Illumina RNA sequencing. We extracted total RNA using miRNeasy Mini Kit (Qiagen, Cat# 217004) from ten sexed pupae: WT ♀, WT♂, transheterozygous *pgSIT*^*♂’*^s, and *pgSIT*^*♀*^harboring all three transgenes Cas9/+; *gRNA*^*βTub#7*^ /+; *gRNA*^*myo-fem#1*^ /+ with each genotype in biological triplicate (12 samples total), following the manufacturer’s protocol. DNase treatment was conducted using DNase I, RNase-free (ThermoFisher Scientific, Cat# EN0521), following total RNA extraction. RNA integrity was assessed using the RNA 6000 Pico Kit for Bioanalyzer (Agilent Technologies #5067-1513), and mRNA was isolated from ∼1 μg of total RNA using NEBNext Poly(A) mRNA Magnetic Isolation Module (NEB #E7490). RNA-seq libraries were constructed using the NEBNext Ultra II RNA Library Prep Kit for Illumina (NEB #E7770) following the manufacturer’s protocols. Briefly, mRNA was fragmented to an average size of 200 nt by incubating at 94°C for 15 min in the first strand buffer. cDNA was then synthesized using random primers and ProtoScript II Reverse Transcriptase followed by second strand synthesis using NEB Second Strand Synthesis Enzyme Mix. Resulting DNA fragments were end-repaired, dA tailed, and ligated to NEBNext hairpin adaptors (NEB #E7335). Following ligation, adaptors were converted to the “Y” shape by treating with USER enzyme, and DNA fragments were size selected using Agencourt AMPure XP beads (Beckman Coulter #A63880) to generate fragment sizes between 250 and 350 bp. Adaptor-ligated DNA was PCR amplified followed by AMPure XP bead clean up. Libraries were quantified using a Qubit dsDNA HS Kit (ThermoFisher Scientific #Q32854), and the size distribution was confirmed using a High Sensitivity DNA Kit for Bioanalyzer (Agilent Technologies #5067-4626). Libraries were sequenced on an Illumina HiSeq2500 in single read mode with the read length of 50 nt and sequencing depth of 20 million reads per library. Base calls were performed with RTA 1.18.64 followed by conversion to FASTQ with bcl2fastq 1.8.4. The reads were mapped to the AaegL5.0 (GCF_002204515.2) genome supplemented with OA-874PA, OA-1067A1, and OA-1067K sequences using STAR. On average, ∼97.5% of the reads were mapped (**Table S11**). Gene expression was then quantified using featureCounts against the annotation release 101 GTF downloaded from NCBI (GCF_002204515.2_AaegL5.0_genomic.gtf). TPM values were calculated from counts produced by featureCounts and combined (**Table S12**).

PCA and hierarchical clustering of the data show that the samples generally behaved as expected in clustering by sex and genotype (**Fig. S10**). DESeq2 was then used to perform differential expression analyses between pgSIT vs WT samples within each sex (**Fig. S10, Table S13, S14**), and a two-factor design consistently showed what changed in response to the genotype in both sexes (**Table S15**). In a comparison between *pgSIT*^*♀*^and WT ♀, 660 genes were upregulated in *pgSIT*^*♀*^and 392 were downregulated at an adjusted p-value < 0.05. The target gene, AAEL005656, was significantly downregulated in *pgSIT*^*♀*^(**Fig. S10C**). In a comparison between *pgSIT*^*♂’*^s andWT♂ (**Table S13**), 2067 genes were upregulated in *pgSIT*^*♂’*^s and 2722 were downregulated at an adjusted p-value < 0.05. The target gene, AEL019894, was strongly downregulated in *pgSIT*^*♂*^ (**Fig. S10D**). It’s important to note here that the CRISPR/Cas9 pgSIT system disrupts the DNA (not the RNA) so transcription is expected to occur; however, the transcripts produced will encode mutations and should be degraded by nonsense -mediated mRNA decay (NMD) mechanisms. Indeed, these mutant RNA’s can be observed in the IGV (**Fig. S8, S9**). In the two-factor comparison, 1447 genes were upregulated in pgSIT and 2563 were downregulated at an adjusted p-value < 0.05 (**Fig. S10E**). For each DESeq2 comparison, gene ontology enrichments were performed on significantly differentially expressed genes, and these are provided as tabs in the corresponding tables (**Table S13-S15**). All Illumina RNA sequencing data has been deposited to the NCBI sequence read archive (SRA) under BioProject ID is PRJNA699282 with accession numbers SRR13620773-SRR13620784.

### Amplicon Sequencing of Target Loci

To sample a variety of molecular changes at the gRNA target sites (*myo-fem* and *βTub*), we used the Amplicon-EZ service by Genewiz® and followed the Genewiz® guidelines for sample preparation. Genomic DNA from 50 WT and 50 pgSIT sexed pupae (25♀+ 25♂) were extracted separately using DNeasy Blood and Tissue Kit (Qiagen, Cat No./ID: 69506) following the manufacturer’s protocols. Primers with Illumina adapters (**Table S18**) were used to PCR amplify the genomic DNA. PCR products were purified using the Zymoclean Gel DNA Recovery Kit (Zymo Research, Cat No./ID: D4007). Roughly 50,000 one-directional reads were generated by Genewiz® and uploaded to Galaxy.org for analysis. Quality control for the reads was performed using FASTQC. Sequence data were then paired and aligned against the *myo-fem* or *βTub* sequence using Map with BWA-MEM under “Simple Illumina mode”. Sequence variants were detected using FreeBayes, with parameter selection level set to “simple diploid calling.” The amplicon sequencing data has been provided as **File S1**.

### Prior mating with *pgSIT*^*♂*^’s suppress ♀fertility

To determine whether prior matings with *pgSIT*^*♂*^ could reduce ♀fertility, we initiated 15 cages each consisting of 250 mature (4–5 days old) *pgSIT*^*♂*^ combined with 50 mature (4–5 days old) WT virgin ♀. We allowed the *pgSIT*^*♂*^’s to mate with these ♀’s for a limited period of time (including: 2, 6, 12, 24, and 48 hrs; 3 replicate cages each). Cages were shaken every 3 minutes for the first half hour to increase mating opportunities. Following these time periods, all ♀’s were removed and transferred to new cages along with 250 WT mature ♂’s, cages were again shaken every 3 minutes for the first half hour to increase mating opportunities and left to mate for an additional 2 days. The ♀’s were then blood fed, and each blood fed ♀was individually transferred to a single narrow Polystyrene vial (Genesee Scientific Cat# 32-116), and eggs were collected and hatched for fertility determination. Following this, non-fertile ♀’s were then placed back into cages along with the original WT ♂’s, plus an additional 50 mature WT ♂’s, for another chance to produce progeny. This was repeated for up to five gonotrophic cycles. As controls, cages with 250 WT ♂’s and 50 WT ♀’s, or 50 unmated blood fed WT ♀’s with no ♂’s added, or 50 unmated blood fed WT ♀’s with 250 WT ♂ adults were also set up (**Table S8**).

### Life table parameters

Life table parameters were assessed by comparing WT, homozygous *gRNA*^*βTub*+*myo-fem*^, homozygous *Cas9*, and transheterozygous pgSIT (*gRNA*^*βTub*+*myo-fem*^/+; Cas9/+) generated with Cas9 inherited from either the mother (maternal Cas9) or father (paternal Cas9). Larva/pupae development times were recorded as the number of days from hatched larvae to pupae and then to adults. One hundred larvae from each line were placed in separate larval rearing containers (Sterilite, 34.6 cm ×21 cm ×12.4 cm, USA), each with 3 liters of deionized water, and fed once a day. Larvae were counted twice daily until pupation, and then the date of pupation and emergence were recorded. Larval to pupae development time was calculated for each sex. Pupae were transferred to plastic cups (Karat, C-KC9) with 100 ml of water, and survivors were recorded until adulthood. ANOVA and Tukey post-hoc tests were performed to compare differences in larval and pupal development among all groups.

For measuring ♂ /♀longevity, we tested the variation in ♂ and ♀longevity among different lines using two methods: (i) released along with WT of the opposite sex or (ii) without WT of the opposite sex. (i) One hundred WT, homozygous *gRNA*^*βTub*+*myo-fem*^, homozygous Cas9 newly eclosed adult mosquitoes (fifty ♂’s and fifty ♀’s) were maintained in a cage; fifty newly-eclosed pgSIT ♂’s (maternal cas9) and fifty newly-eclosed pgSIT ♂’s (paternal cas9) were caged with fifty newly-eclosed WT ♀’s; and finally, fifty newly-eclosed pgSIT ♀’s (maternal cas9) and fifty newly-eclosed pgSIT ♀’s (paternal cas9) were caged with fifty newly-eclosed WT ♂’s. (ii) Fifty\ ♂’s or ♀’s from each line were released into a cage separately without the opposite sex. Adults were provided with 10% sucrose and monitored daily for survival until all mosquitoes had died (3 replicates).

For measuring ♀fecundity and fertility, ♀’s (n = 50) and ♂’s (n = 50) three days post-emergence raised under the same standardized larval conditions were placed into a cage and allowed to mate for 2 days. ♀mosquitoes were blood fed until fully engorged and were individually transferred into plastic vials with oviposition substrate. Eggs were stored in the insectary for 4 days to allow full embryonic development and then were hatched in a vacuum chamber. Fecundity was calculated as the number of eggs laid per ♀, and fertility was calculated as the percentage of eggs hatched per ♀. An analysis of variance (ANOVA) and a Tukey post-hoc test were performed to compare differences in fecundity and fertility among all groups.

♂ mating capacity (how many ♀’s can be mated by one mature ♂) was measured as follows. Fifteen mature WT ♀’s were caged with 1 mature ♂ of each genotype for 24 hours (1♂:15♀ratio). After 24 hours, the single ♂ was removed from all cages. Two days after the single ♂ was removed, 75 WT ♂’s were added to each cage that previously had a pgSIT ♂ (5♂:1♀ratio). Blood meals were provided, and each blood fed ♀was individually transferred to a single vial for egg collection. The fecundity and fertility of each ♀was determined. The mating capacity was calculated as the total number of ♀’s - total number of fertile ♀’s. The mating capacity of WT, homozygous *gRNA*^*βTub*+*myo-fem*^, and homozygous Cas9 ♂ was equal to the number of fertile ♀’s. All statistical analyses were performed using GraphPad Prism software (GraphPad Software, La Jolla, California, USA). P values > 0.05 were considered not significant.

### Flight activity quantification

Mosquitoes were reared at 28°C, 80% relative humidity under a 12:12 hr light:dark regime, and measurements of flight activity were performed using a *Drosophila* Activity Monitoring (DAM) System (TriKinetics, LAM25) using large tubes designed for mosquitoes (TriKinetics, PGT 25 x 125 mm Pyrex Glass). Individual 4–7 day-old, non-blood fed virgin ♀and non-mated ♂ mosquitoes were introduced into the monitoring tubes, which contained 10% sucrose (Sigma, Cat. S0389) at both ends of the tube as the food source. The DAM System was positioned vertically during the assays. Flight activity was measured over a period of 24 hrs by automatically calculating the number of times that mosquitoes passed through the infrared beam in the center of the tubes. The walls of the monitoring tubes were coated with Sigmacote (Sigma, Cat. SL2) to inhibit mosquitoes from walking upward. For preparing the wingless mosquitoes, the animals were anesthetized on ice, and the wings were removed using Vannas Scissor (World Precision Instruments, Cat. 14003). The wingless mosquitoes were allowed to recover for 12 hrs before recording. Mosquitoes were manually checked after flight activity recording to ensure survival. Data acquisition was performed using the DAMSystem (TriKinetics) (**Fig. 2D, Video S5, Table S6**).

### Sound attraction assay

The sound attraction assay was performed in a chamber with a temperature of 28°C and humidity of 80%. Seven-day old ♂’s were sex separated after the pupae stage. The day before testing, 30–40 ♂’s were transferred by mouth aspiration to a 15-cm^3^ mesh cage with a 10% sucrose bottle. ♂ mosquitoes were allowed to recover in the cage under a 12 hr:12 hr light:dark regime for 24 hrs. For each trial, a 10-second 600 Hz sine tone was applied on one side of the cage as a mating behavior lure, mimicking ♀flight tones. The number of mosquitoes landing on the mesh area around the speaker box(10 cm^*2*^)was quantified at 5-second intervals throughout the stimulus. The average percent of mosquitoes landing around the speaker area out of the total cage post-sound presentation was calculated (**Fig. 2E, Video S6, Table S7**). Heatmaps were generated using Noldus Ethovision XT.

### Multigenerational population cage trials

To perform multigenerational population cage trials, two strategies were employed: (i) release of eggs; (ii) release mature adults (**Fig. 3, Table S16**). Cage trials were carried out using discrete non-overlapping generations. For the first release of eggs strategy (i), WT eggs and pgSIT eggs were hatched together using the following ratios of 1:1 (100:100), 1:5 (100:500), 1:10 (100:1000), 1:20 (100:2000), and 1:40 (100:4000), and three biological replicates for each ratio (15 cages total). All eggs were hatched simultaneously, then separated into multiple plastic containers (Sterilite, 34.6 cm ×21 cm ×12.4 cm, USA). Roughly 400 larvae were reared in each container using standard conditions with 3 liters of deionized water and were allowed to develop into pupae. Pupae were placed in plastic cups (Karat, C-KC9) with ∼100 ml of water (∼150 pupae per cup) and transferred to large cages (BugDorm, 60cm ×60cm ×60cm) to eclose. All adults were allowed to mate for ∼5–7 days. ♀’s were blood fed, and the eggs were collected. Eggs were counted and stored for ∼4 days to allow full embryonic development, then 100 eggs were selected randomly and mixed with pgSIT eggs with ratios of 1:1 (100:100), 1:5 (100:500), 1:10 (100:1000), 1:20 (100:2000), and 1:40 (100:4000) to seed for the following generation, and this procedure continued for all subsequent generations. The remaining eggs were hatched to measure hatching rates and to screen for the possible presence of transformation markers. The hatching rate was estimated by dividing the number of hatched eggs by the total number of eggs.

For the release of mature adults strategy (ii), 3–4-days-old mature WT adult ♂’s were released along with mature (3–4 days old) pgSIT adult ♂’s at release ratios: 1:1 (50:50), 1:5 (50:250), 1:10 (50:500), 1:20 (50:1000), and 1:40 (50:2000), with three biological replicates for each release ratio (15 cages total). One hour later, 50 mature (3–4 days old) WT adult ♀’s were released into each cage. All adults were allowed to mate for 2 days. ♀’s were then blood fed and eggs were collected. Eggs were counted and stored for four days to allow full embryonic development. Then, 100 eggs were randomly selected, hatched, and reared to the pupal stage, and the pupae were separated into ♂ and ♀groups and transferred to separate cages. Three days post eclosion, 50 (1:1), 250 (1:5), 500 (1:10), 1000 (1:20), and 2000 (1:40) age-matched pgSIT mature ♂ adults were caged with these mature ♂’s from 100 selected eggs. One hour later, mature ♀’s from 100 selected eggs were transferred into each cage. All adults were allowed to mate for 2 days. ♀’s were blood fed, and eggs were collected. Eggs were counted and stored for 4 days to allow full embryonic development. The remaining eggs were hatched to measure hatching rates and to screen for the possible presence of transformation markers. The hatching rate was estimated by dividing the number of hatched eggs by the total number of eggs. This procedure continued for all subsequent generations.

### Mathematical modeling

To model the expected performance of pgSIT at suppressing and eliminating local *Ae. aegypti* populations, we used the MGDrivE simulation framework (*19*). This framework models the egg, larval, pupal, and adult mosquito life stages with overlapping generations, larval mortality increasing with larval density, and a mating structure in which females retain the genetic material of the adult ♂ with whom they mate for the duration of their adult lifespan. The inheritance pattern of the *pgSIT* system was modeled within the inheritance module of MGDrivE, along with impacts on adult lifespan, male mating competitiveness, and pupatory success. We distributed *Ae. aegypti* populations according to human structures sourced from OpenStreetMap on the basis that *Ae. aegypti* is anthropophilic. Each human structure was assumed to have an equilibrium population of 16 adult *Ae. aegypti*, producing an equilibrium island population of 992. We implemented the stochastic version of the MGDrivE framework to capture random effects at low population sizes and the potential for population elimination. Weekly releases of up to 400 *pgSIT* eggs were simulated in all human structures of Onetahi over a period of 10-24 weeks. 200 repetitions were carried out for each parameter set, and mosquito genotype trajectories, along with the proportion of simulations that led to local population elimination, were recorded. Complete model and intervention parameters are listed in **Table S17**.

## Acknowledgments

We thank Judy Ishikawa for helping with mosquito husbandry. This work was supported by funding from a DARPA Safe Genes Program Grant (HR0011-17-2-0047) and an NIH award (R01AI151004) awarded to O.S.A. and an NIH award (R56-AI153334) to C.M. The views, opinions, and/or findings expressed are those of the authors and should not be interpreted as representing the official views or policies of the U.S. government.

## Author Contributions

O.S.A and M.L. conceptualized and designed experiments; T.Y., J.L., L.A., S.G., performed molecular analyses; M.L., J.R.E., T.W., H.L., M.B. performed genetic experiments; Y.Z, Y.W., N.D., J.C, C.M. performed behavioral experiments; J.B., H.M.S.C, J.M.M performed mathematical modeling; I.A. performed bioinformatics: O.S.A, M.L., N.P.K, R.R., analyzed and compiled the data. All authors contributed to writing and approved the final manuscript.

## Ethical conduct of research

All animals were handled in accordance with the Guide for the Care and Use of Laboratory Animals as recommended by the National Institutes of Health and approved by the UCSD Institutional Animal Care and Use Committee (IACUC, Animal Use Protocol #S17187) and UCSD Biological Use Authorization (BUA #R2401).

---

**Figure S1.**
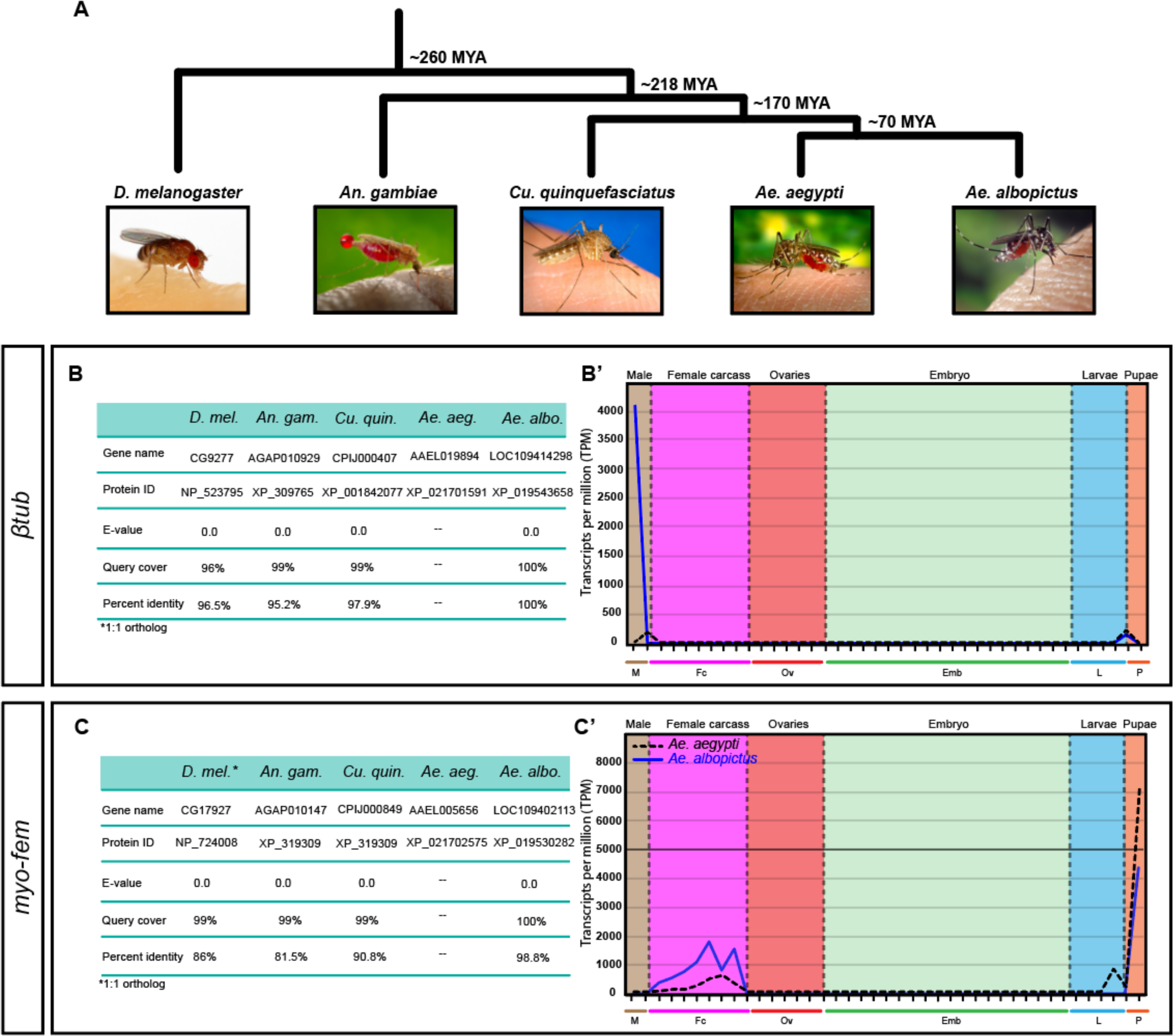
Conservation of target genes in Diptera. (**A**) Phylogenetic tree of Dipteran insects, including *Drosophila melanogaster (D. melanogaster and D. mel*.*), Anopheles gambiae (An. gambiae* and *An. gam*.*), Culex quinquefasciatus (Cu. quinquefasciatus* and *Cu. quin*.*), Aedes aegypti (Ae. aegypti* and *Ae. aeg)*, and *Aedes albopictus (Ae. albopictus* and *Ae. albo)*, with evolutionary distance measured by million years ago (MYA). All images are free to use and were downloaded from wiki commons, except *Cu. quin*., which was downloaded from Pixnio.com. (**B**) *βTub* orthologs in Dipteran species with highest similarity to the *Ae. aegypti βTub* gene at the amino acid level. (B’) RNA expression levels of the *βTub* gene throughout development. Extremely high expression of *βTub* is seen in *Ae. albopictus* testes samples (refer to **Table S1** for TPM data of *βTub* expression across development). Lower gene expression is observed in both ♂ carcass and ♂ pupae samples. (**C**) *myo-fem* orthologs in Dipteran species with the highest similarity to the *Ae. aegypti myo-fem* gene at the amino acid level. (C’) RNA expression levels of the myosin gene across development in both *Ae. aegypti* and *Ae. albopictus* mosquito samples. The *myo-fem* gene is highly expressed in the pupal samples of both mosquito species (refer to **Table S1** for TPM data of *myo-fem* expression across development). **(B’ and C’)** RNA-Seq expression levels of *myo-fem* and *βTub* genes in *Ae. aegypti* and *Ae. albopictus* mosquito samples using available data (*11, 12*). Major stages of development are color coded, where brown represents ♂ testes and ♂ carcass samples, pink represents ♀carcass samples, red represents ovary samples, green represents embryogenesis, blue represents larval samples, and orange represents pupae samples. Black dotted lines represent *Ae. aegypti*, and solid blue lines represent *Ae. albopictus*. The major developmental groups are indicated by color bars and are organized left to right, as follows: M (brown, ♂ testes, ♂ carcass), Fc (purple, NBF ♀Carcass, and multiple time points PBM: 12, 24, 36, 48, 60, and 72 hr), Ov (red, NBF ovaries, and multiple ovarian time points PBM: 12, 24, 36, 48, 60, and 72 hr), Emb (green, embryo, 0–2, 2–4, 4–8, 8–12, 12– 16, 16–20, 20–24, 24–28, 28–32, 32–36, 36–40, 40–44, 44–48, 48–52, 52–56, 56–60, 60–64, 64–68, 68–72, and 72–76 hr embryos), L (light blue, larvae, 1st, 2nd, 3rd, and 4th instar larvae stages), and P (light orange, ♂ and ♀pupae). *1:1 orthologs.

**Figure S2.**
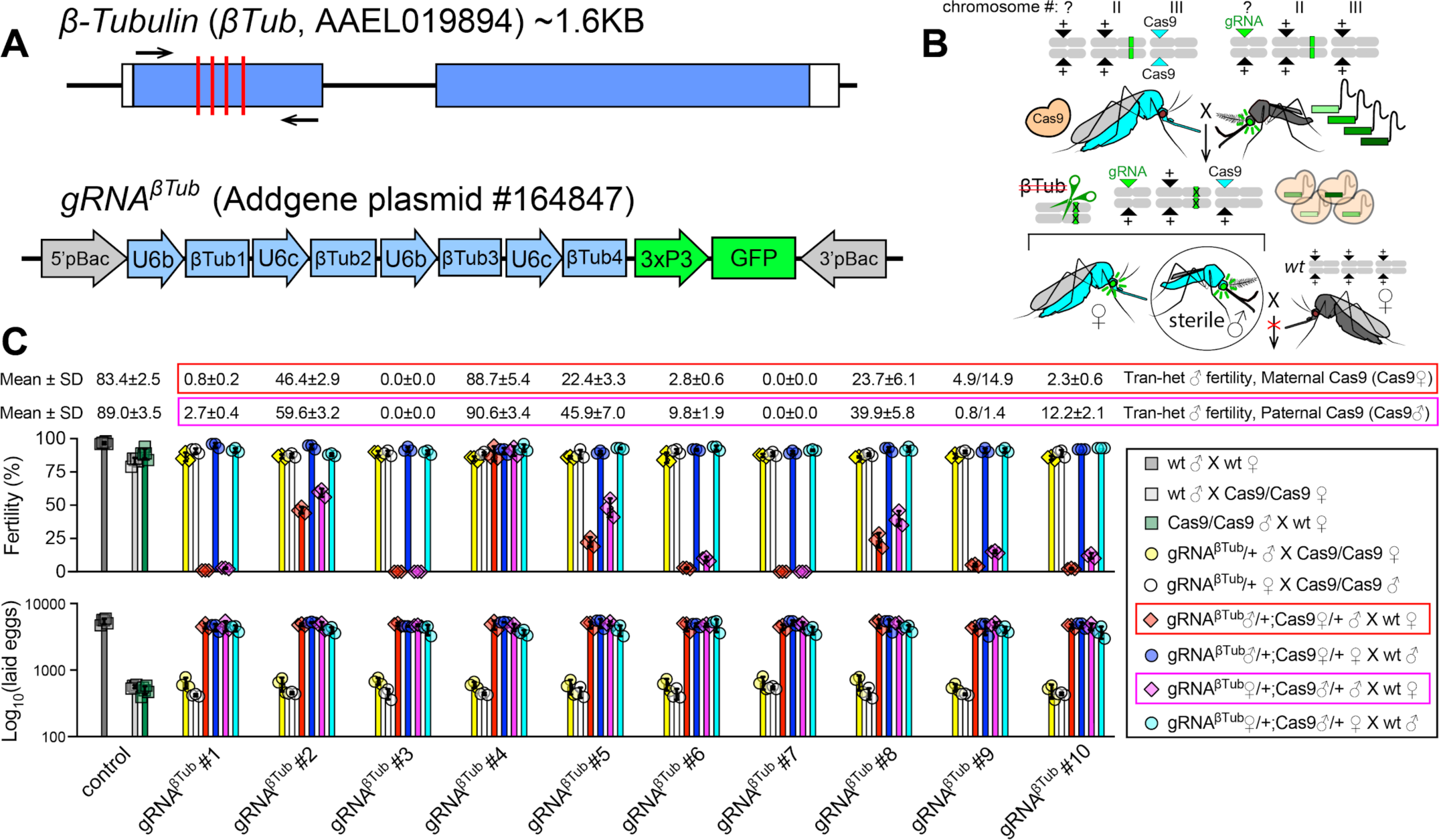
Assessment of independent *gRNA*^*βTub*^ lines. (**A**) Schematic of the *Ae. aegypti β-Tubulin* (*βTub*) locus and the *gRNA*^*βTub*^ construct used to generate 10 transgenic *gRNA*^*βTub*^ lines. The *gRNA*^*βTub*^ construct harbors four independent gRNAs targeting different sequences in the 1st coding exon of *βTub* (red lines) and a *3xP3-GFP* marker gene. (**B**) A schematic of the genetic cross between the homozygous *Cas9* ♀’s and heterozygous *gRNA*^*βTub*^/+ ♂’s generating the transheterozygous progeny inheriting maternal Cas9. To assess the efficiency of *βTub* disruption in F_1_ transheterozygous progeny, this cross was set up reciprocally between the homozygous *Cas9* line and each of ten different insertion lines of *gRNA*^*βTub*^. The generated F_1_ transheterozygous ♂’s and ♀’s were crossed to WT mosquitoes of the opposite sex, and their fertility and fecundity, as an average number of laid eggs, were compared to those of F_0_ parents, homozygous Cas9, and WT mosquitoes. (**C**) The bar plots show biological replicates and means ± SDs for fertility and fecundity (Log_10_[laid eggs]) of tested groups (**Table S3**). Different insertion lines of the same *gRNA*^*βTub*^ construct in conjunction with *Cas9* induced a range of infertility in transheterozygous ♂’s from 90.6 ± 3.4% to 0%. Two lines, *gRNA*^*βTub*^*#3* and *gRNA*^*βTub*^*#7*, independently induced the robust sterility of transheterozygous ♂’s where they harbored maternal or paternal Cas9: *gRNA*^*βTub*^♂/+; Cas9♀/+ or *gRNA*^*βTub*^♀/+; Cas9♂/+, respectively. The *gRNA*^*βTub*^*#7* was the easiest to score due to its brighter expression of the *3xP3-GFP* marker. Therefore, it was used for further analysis and genetically combined with the best *gRNA*^*myo-fem*^ line. Statistical significance was estimated using a two-sided Student’s *t* test with unequal variance. (*p* ≥ 0.05^ns^, *p* < 0.05*, *p* < 0.01**, and *p* < 0.001***).

**Figure S3.**
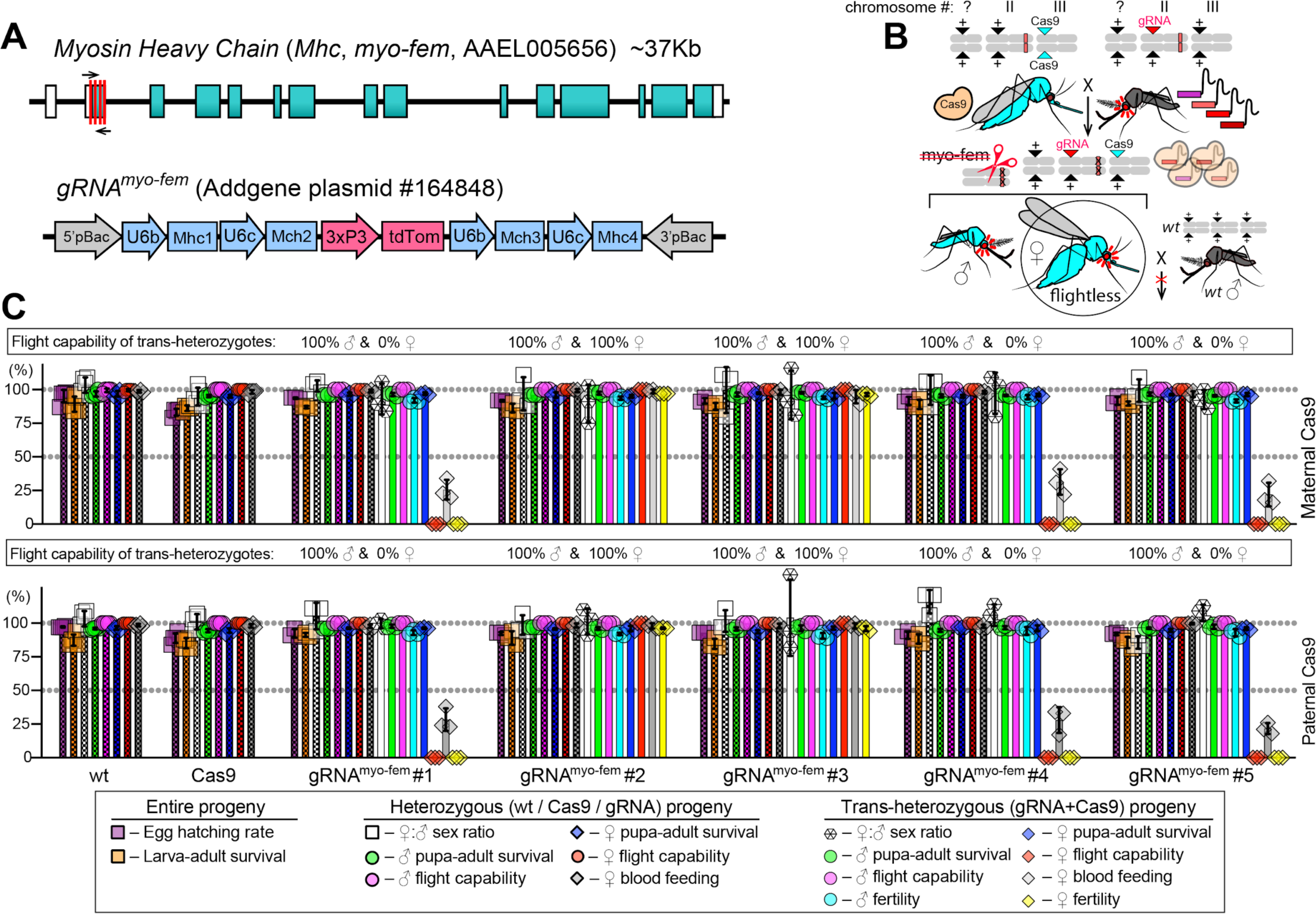
Assessment of independent *gRNA*^*myo-fem*^ lines. (**A**) Schematic of the *Ae. aegypti Myosin Heavy chain* (*Mhc, myo-fem*) locus and the *gRNA*^*myo-fem*^ genetic construct used to generate five transgenic *gRNA*^*myo-fem*^ lines. The *gRNA*^*myo-fem*^ construct harbors four independent gRNAs targeting different sequences in the 1st exon of *myo-fem* (red lines) and a *3xP3-tdTomato* marker gene. (**B**) A schematic of the genetic cross between the homozygous *Cas9* ♀’s and heterozygous *gRNA*^*myo-fem*^/+ ♂’s generating the transheterozygous progeny inheriting maternal Cas9. To assess the efficiency of *myo-fem* disruption in F_1_ transheterozygous progeny, this cross was set up in both reciprocal directions between the homozygous *Cas9* line and each of five different insertion lines of *gRNA*^*myo-fem*^. We scored the flight ability, pupal lethality, blood feeding, and fertility in generated F_1_ transheterozygous ♂’s and ♀’s and compared to WT and Cas9 mosquitoes. (**C**) The bar plots show biological replicates and means ± SDs for assessed characteristics (**Table S3**). In the presence of *Cas9*, three out of five *gRNA*^*βTub*^ lines induced the *myo-fem* disruption that resulted in the complete ♀-specific flightlessness, while transheterozygous ♂’s were able to fly. The ♀inability to fly affected their blood feeding, mating, and survival, rendering transheterozygous ♀’s infertile. The *gRNA*^*myo-fem*^*#1* was the easiest to score due to its brighter expression of the *3xP3-tdTomato* marker. Therefore, it was used for further analysis and genetically combined with the *gRNA*^*βTub*^*#7* line. Statistical significance was estimated using a two-sided Student’s *t* test with unequal variance. (*p* ≥ 0.05^ns^, *p* < 0.05*, *p* < 0.01**, and *p* < 0.001***).

**Figure S4.**
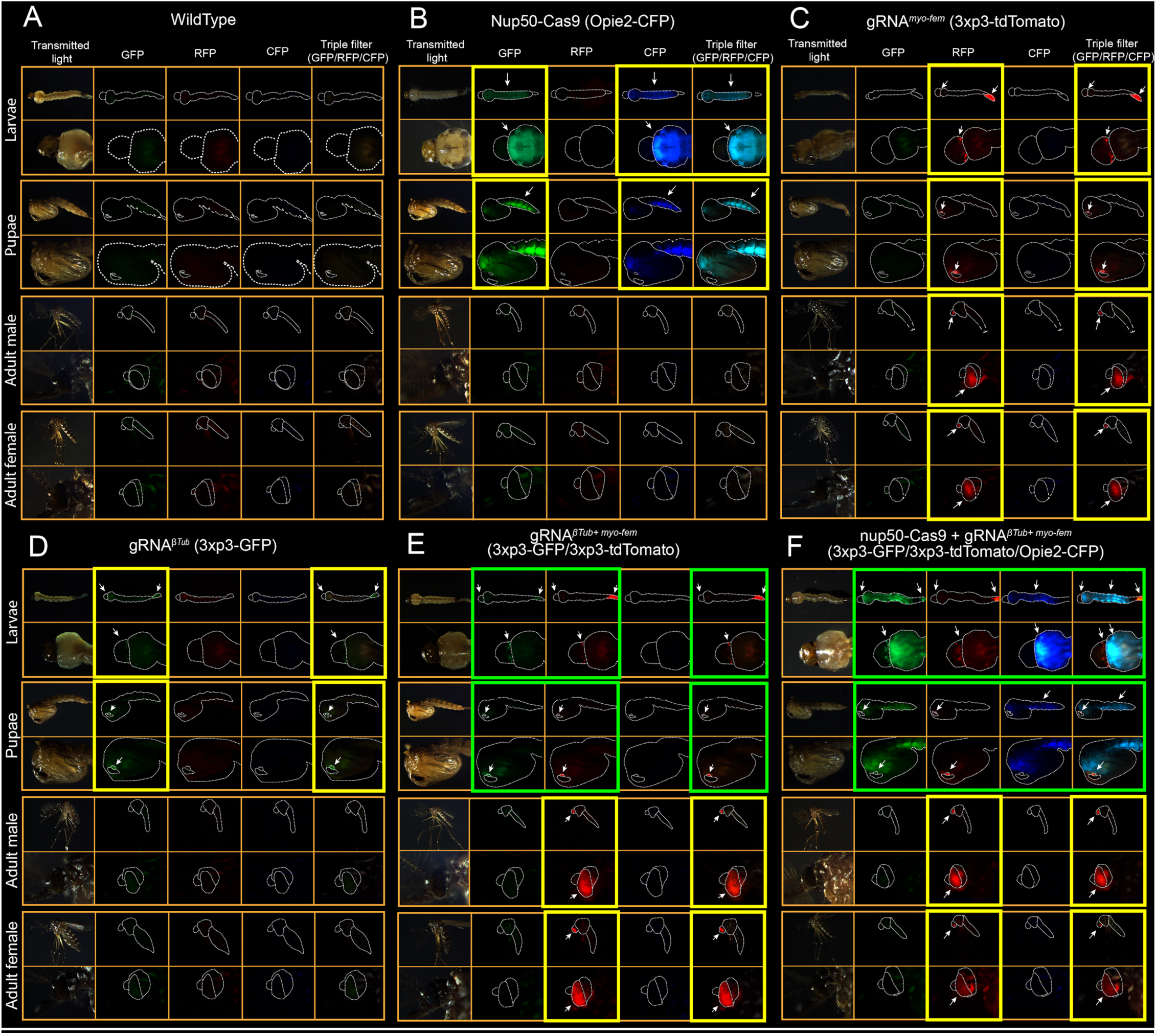
Transmitted light and fluorescent images of mosquito life stages of strains used in this study. For each strain, multiple life stages are shown, including larvae (instar 4), pupae, and adult ♂’s and ♀’s. All stages are imaged using transmitted white light, GFP filter (Leica Part #10447408; ET470/40x, ET525/50m wavelengths), RFP filter (Leica Part #10450195; ET560/40x, ET630/75m wavelengths), CFP filter (Leica Part #10447409; ET436/20x, ET480/40m wavelengths) and a triple filter for GFP/RFP/CFP (Leica Part #10450611; ET434.5/21, 501.5/19, 574.5/23, ET469.5/25, 536.5/29, 635.5/69 wavelengths) using a Leica M165FC fluorescent stereomicroscope. **(A)** WT mosquitoes. **(B**) Cas9 marked with Opie2-CFP. **(C)** gRNA^*myo-fem*^ marked with 3xp3-tdTomato. **(D)** gRNA^*βTub*^ marked with 3xp3-GFP. **(E)** gRNA^*βTub+myo-fem*^ marked with 3xp3-GFP and 3xp3-tdTomato. **(F)** Cas9 + gRNA^*βTub+myo-fem*^ marked with Opie2-CFP, 3xp3-GFP, and 3xp3-tdTomato. Arrows indicate points of focus where markers can easily be distinguished. Yellow box indicates the life stage in which the transgene can be reliably distinguished. Green box indicates the life stage in which mosquitoes harboring multiple transgenes can be reliably distinguished in E,F.

**Figure S5.**
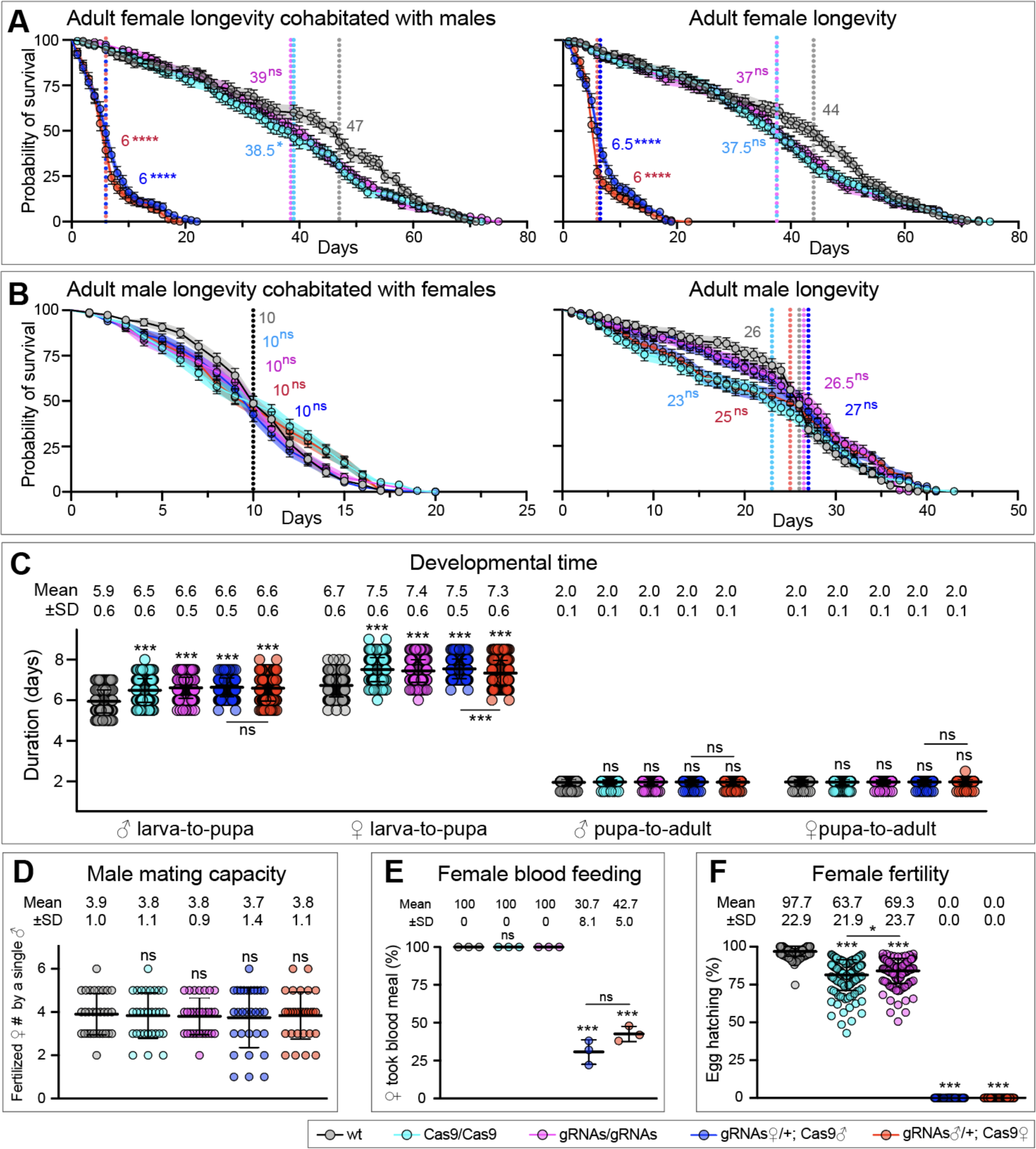
Fitness of transheterozygous pgSIT mosquitoes in comparison with WT and parental lines. Survival plots of adult ♀’s either cohabitated with ♂’s or not (**A**), and adult ♂’s either cohabitated with ♀’s or not (**B**). Survival means ± standard errors (SE) over days following adult eclosion are plotted. Vertical lines and values present median survivals for each tested group. Survival curves were compared to the curve for WT’s of the corresponding sex. The departure significance was assessed with the Log-rank (Mantel-Cox) and Gehan-Breslow-Wilcoxon texts and is indicated above median values. The flightlessness of *pgSIT*^♀^’s affected their survival and drastically reduced their longevity even in the laboratory setting. Notably, the longevity of *pgSIT*^*♂*^’s was not significantly affected. (**C**) Larva-to-pupa and pupa-to-adult developmental times were measured in 150 ♀’s and ♂’s. (**D**) Plots of mating capacities for adult ♂’s of each genotype. (**E**) Plots of blood feeding rates of adult ♀’s of each genotype. **(F)** Plots of fertility of adult ♀’s of each genotype. Point plots in panels **C, D, E**, and **F** show mean ± standard deviation (SD). Statistical significance was estimated using a two-sided Student’s *t* test with unequal variance. (*p* ≥ 0.05^ns^, *p* < 0.05*, *p* < 0.01**, *p* < 0.001***, and *p* < 0.0001****).

**Figure S6.**
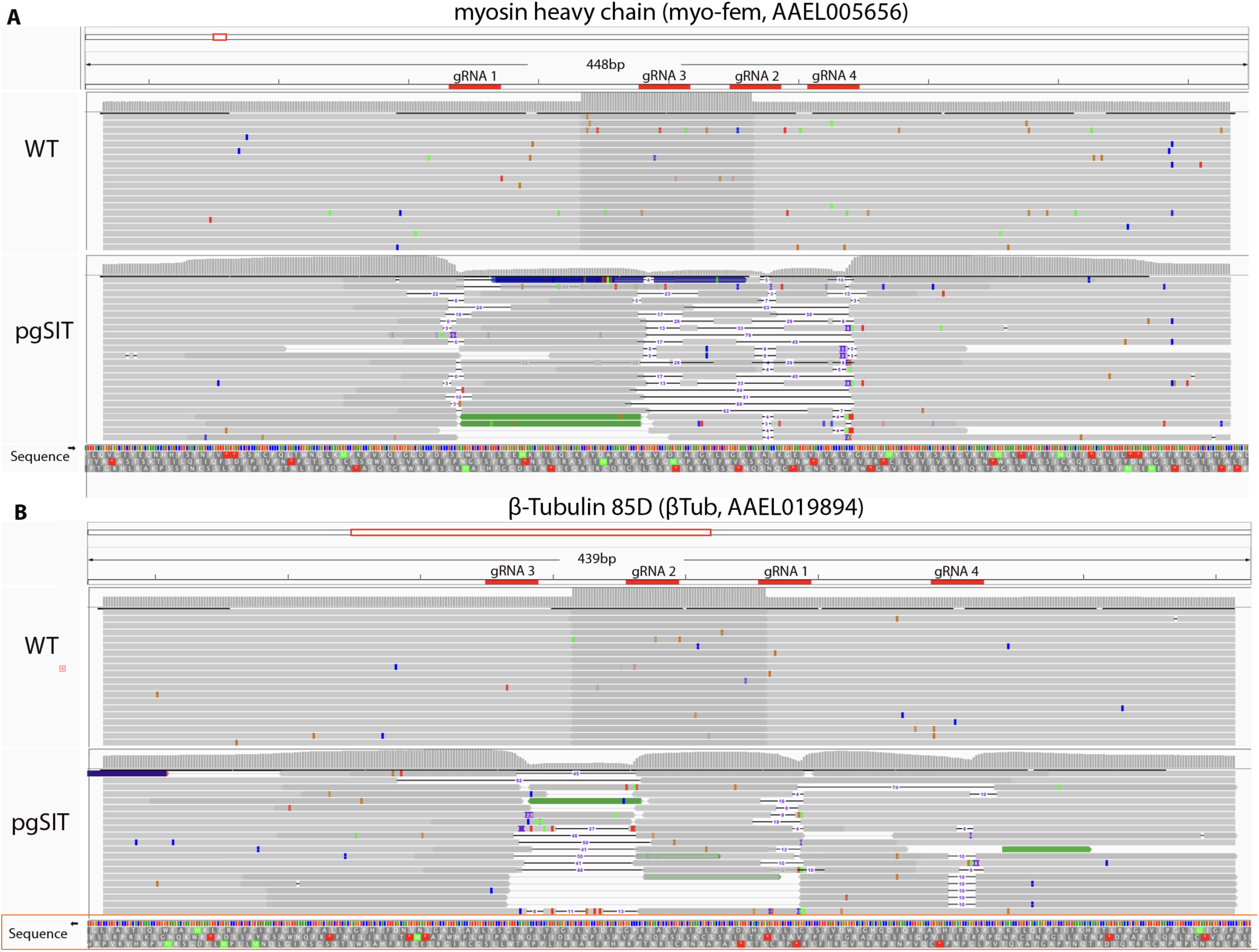
Illumina NGS-based amplicon sequencing results representing *myo-fem* and *βTub* knockout in pgSIT mosquitoes. A zoomed in genome browser snapshot depicting amplicon sequencing based insertions/deletions (indels) at each gRNA target site of: (**A**) *myo-fem* exon 1 in WT individuals (25♀+ 25♂) and pgSIT individuals (25♀+ 25♂). (**B**) *βTub* exon 1 in WT individuals (25♀+ 25♂) and pgSIT individuals (25♀+ 25♂).

**Figure S7.**
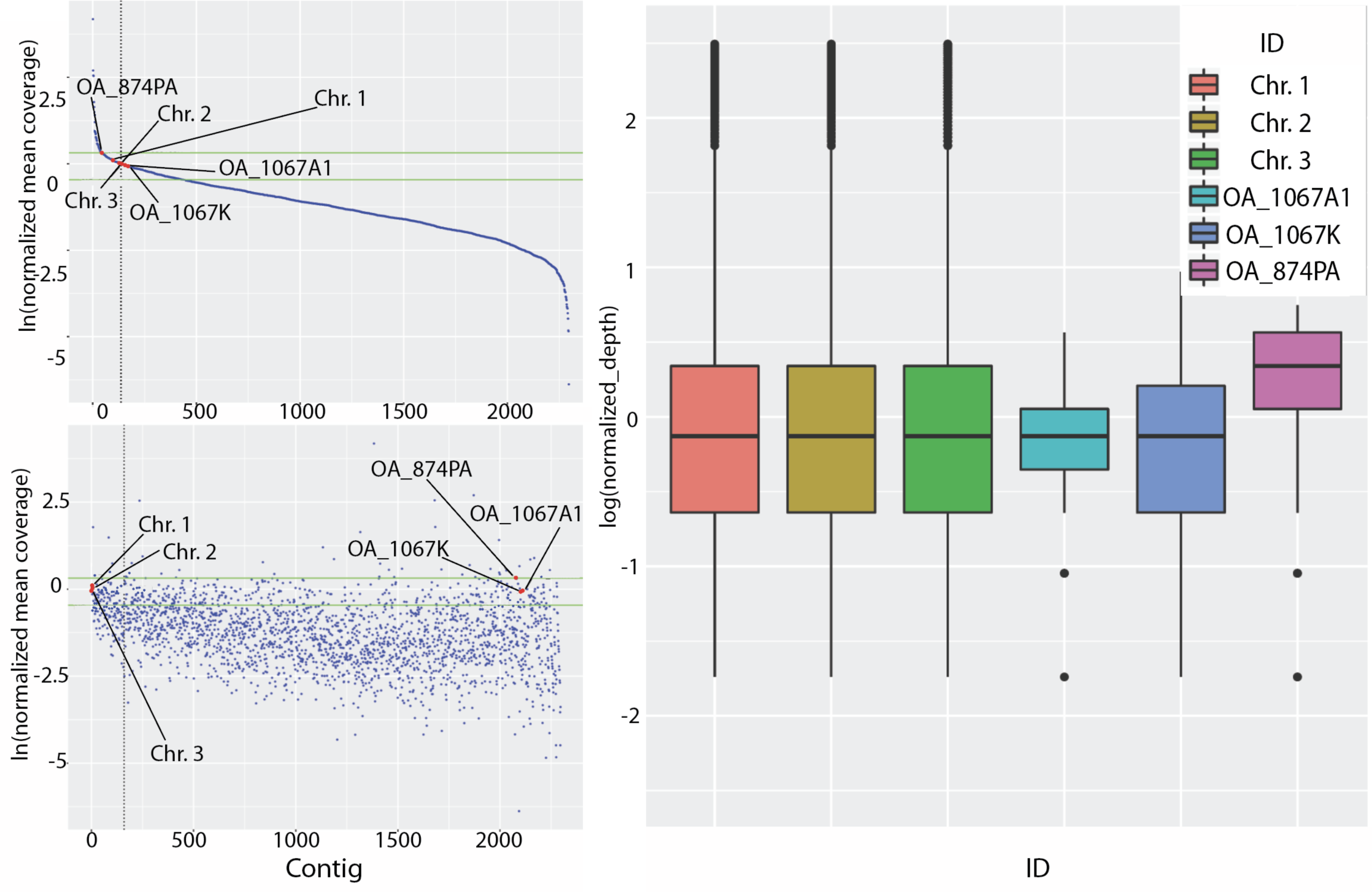
Determination of transgene copy number using Oxford Nanopore genome sequencing. The normalized mean coverage for all the contigs in the genome with the exception of mtDNA, which has a coverage of 6563. These are plotted (blue) by sorting the order of the contigs as either (A) a mean coverage to produce a smooth line or (B) by contig size. The green horizontal lines correspond to the standard deviation. (C) A box plot depicting the coverage distributions of the three chromosomes and the three trangenes. Data associated with this figure can be found in **Tables S9-S10**.

**Figure S8.**
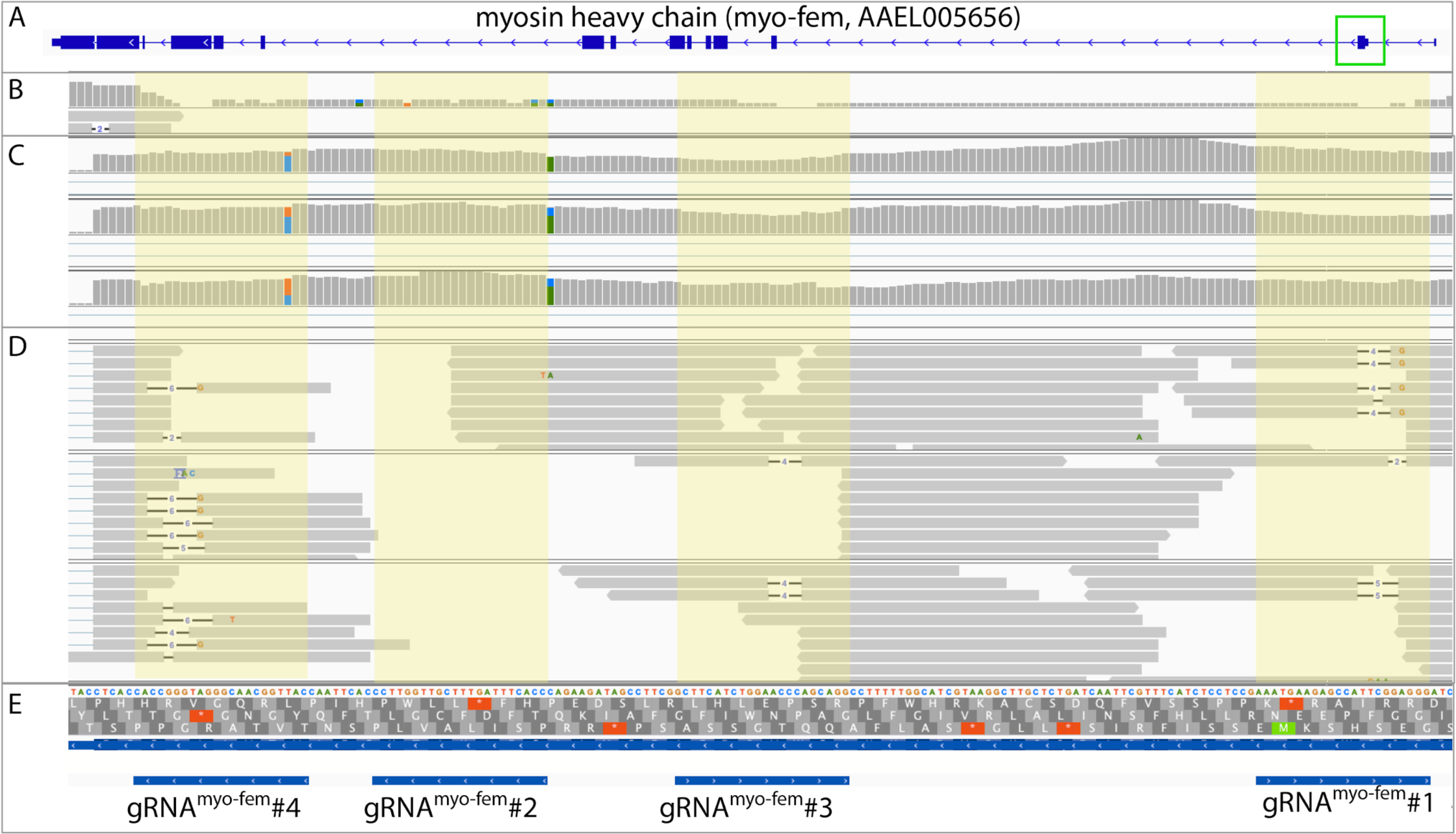
Integrated genome browser snapshot depicting pgSIT sequencing results for *myo-fem*. **(A)** A zoomed out image of the *myo-fem* gene structure with a green box highlighting exon 1 targeted by the gRNAs. **(B-E)** A zoomed in genome browser snapshot of exon 1 depicting: **(B)** Oxford nanopore sequencing results depicting CRISPR/Cas9-mediated mutations in the DNA sequence of *myo-fem* exon 1. **(C)** Illumina transcriptome RNA-sequencing results of WT sequences (3 replicates) showing the lack of mutations as compared to **(D)** depicting mutations in the coding sequence of *myo-fem* exon 1 in pgSIT individuals (3 replicates). **(E)** Depicts the precise locations of the four gRNA target sites.

**Figure S9.**
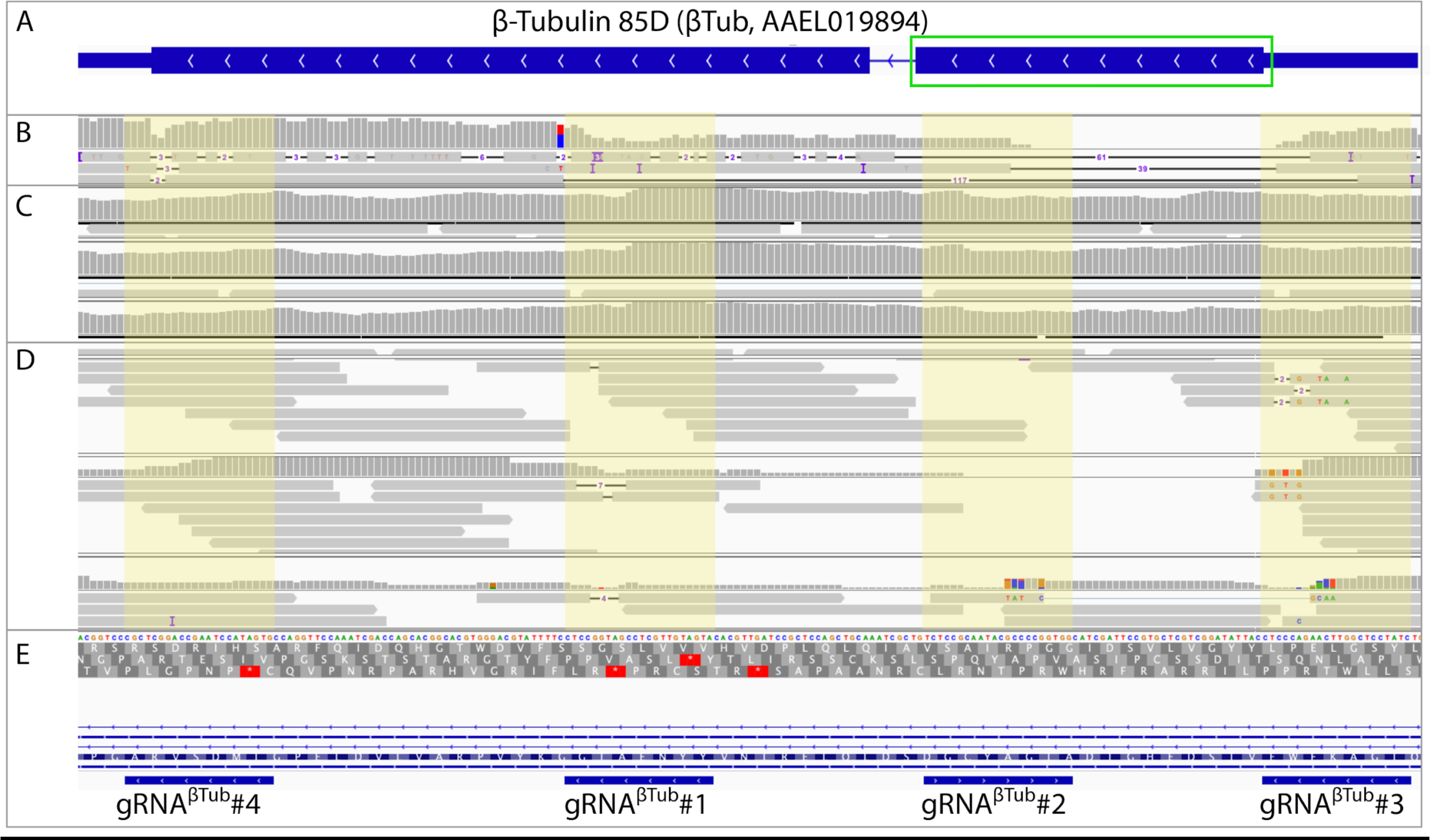
Integrated genome browser snapshot depicting pgSIT sequencing results for *βTub*. A genome browser snapshot of *βTub* depicting: **(A)** A zoomed out image of the *βTub* gene structure with a green box highlighting the exon targeted by gRNAs. **(B-E)** A zoomed in genome browser snapshot of exon-1 depicting: **(B)** Oxford nanopore sequencing results depicting disruptions in the DNA sequence of *βTub* exon 1. **(C)** Illumina transcriptome RNA-sequencing of WT sequences (three replicates) showing the lack of mutations as compared to **(D)** depicting mutations in the coding sequence of *βTub* exon 1 in pgSIT individuals (three replicates). **(E)** Depicts the precise locations of the four gRNA target sites.

**Figure S10.**
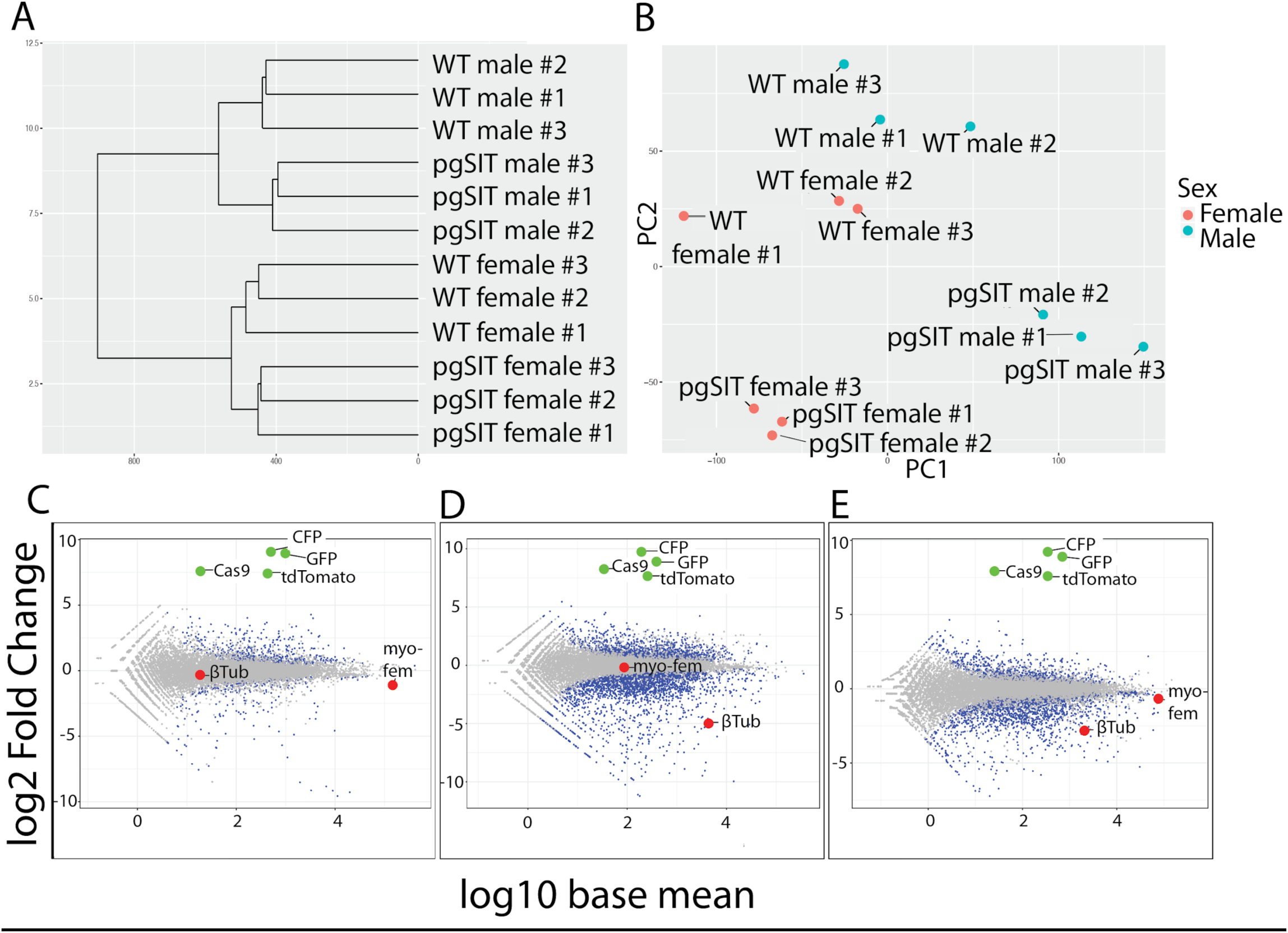
Transcriptional profiling and expression analysis. **(A)** Hierarchical clustering and **(B)** PCA analysis of the 12 samples used for RNA sequencing. **(C-E)** MA-plots showing the differential expression patterns between: **(C)** *pgSIT*^*♀*^vs WT ♀, **(D)** *pgSIT*^*♂’*^s vs WT ♂, **(E)** two-factor *pgSIT* vs WT. Significantly differentially expressed genes are indicated by blue dots (FDR < 0.5), non-significantly differentially expressed genes are indicated by grey dots, target genes are indicated by red dots, transgene encoded genes are indicated by green dots. Data associated with this figure can be found in **Tables S11-15**.

**Figure S11.**
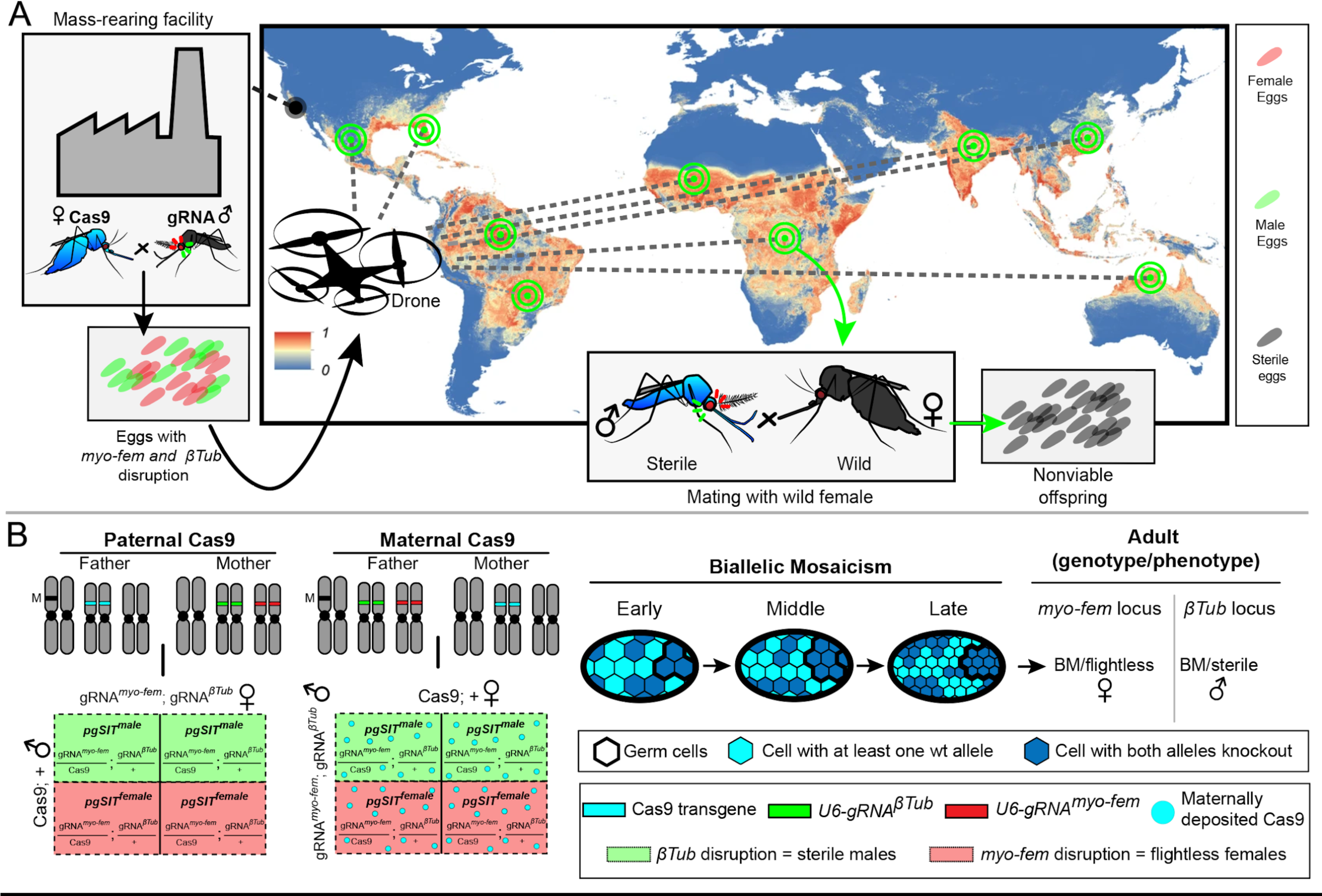
Scaling pgSIT to control populations of mosquitoes and molecular mechanisms. **(A)** A factory produces pgSIT eggs for distribution and release at remote locations worldwide. The global map depicts the probability of occurrence of *Ae. aegypti* (from 0 blue to 1 red) at a spatial resolution of 5 km × 5 km (adopted from (*28*)). **(B)** Punnett squares depict the F1 genotypes derived from bidirectional crosses between homozygous *gRNA*^*βTub*+*myo-fem*^ and *Cas9*. To the right, is a schematic depicting the genetic outcome of a pgSIT cross and the biallelic mosaicism mechanism ensuring F1 flightless ♀and sterile ♂ phenotypes (*9*).

## Supplemental Tables

**Table S1. myo-fem and βTub developmental gene expression data**. *Ae. albopictus* and *Ae. aegypti myo-fem* and *βTub* TPM gene expression across different developmental timepoints

**Table S2. Embryo microinjection and transgenic line generation Table S3. Single gene disruption**

**Table S4. pgSIT cross data**

**Table S5. Life Parameters**. Comparisons of life parameters of different mosquito lines. To evaluate the potential fitness costs associated with the pgSIT components, several life-table parameters such as fecundity, larval development time, ♂ insemination capacity, mating competitiveness, and adult survival rate were measured among WT, homozygous *gRNA*^*βTub*+*myo-fem*^, homozygous Cas9 and heterozygous pgSIT lines (*gRNA*^*βTub*+*myo-fem*^,+/Cas9,+). Compared to WT, homozygous *gRNA*^*βTub*+*myo-fem*^, homozygous Cas9, and *pgSIT*^*♂*^’s that produced from maternal Cas9 or paternal Cas9 had no significant differences in larval and pupal survival rate, larval and pupal development time, ♂ insemination ability, and adult survival (*p* < 0.05). Maternal Cas9 refers to paternal gRNAβTub/+; gRNAmyo-fem/+; maternal Cas9/+, and paternal Cas9 corresponds to maternal gRNAβTub/+; gRNAmyo-fem/+; paternal Cas9/+. Each strain is labeled with a superscript (a-e) and is used to indicate where the significance lies. For example, an ANOVA with a post hoc Tukey’s analysis for ♀fecundity indicates differences are between wildtype vs gRNA, wildtype vs Cas9, and wildtype vs transheterozygotes. **Table S6. pgSIT flight capacity assay**. Flight activities were monitored over a 24-hour period using the DAM system. The counts are the number of times the mosquitoes passed the infrared beam.

**Table S7. Sound attraction assay Table S8. pgSIT**^**♂**^ **sterilize WT ♀’s**

**Table S9**. The mean coverage depth from the Nanopore DNA sequencing for all contigs in the genome (2310) and the three plasmids (OA-1067A1: *gRNA*^*βTub*^; OA-1067K: *gRNA*^*myo-fem*^; and OA-874PA: Nup50-Cas9) as well as normalized coverage based on the global number (**Table S12**). Transgene coverage ranged from 5.1 to 7.6 and normalized coverage ranged from 0.93 to 1.38.

**Table S10**. Nanopore coverage means

**Table S11**. Mapping Stats for RNA sequencing

**Table S12**. RNAseq expression data

**Table S13**. deseq2_liverpool_males_pgSIT_males.annotations

**Table S14**. deseq2_liverpool_females_pgSIT_females.annotations.xlsx

**Table S15**. deseq2_liverpool_pgSIT.annotations

**Table S16. Multigenerational population cage data**

**Table S17**. Parameters used in *Aedes aegypti* population suppression model.

**Table S18. Primer and gRNA sequences**

## Supplemental Videos

**Video S1. Timelapse of *βTub* mutant and WT testes and sperm**. *βTub* mutant and WT testes were imaged at 10X and 63X.

**Video S2. *myo-fem* mutant ♀’s eclosing**. Flightless *myo-fem* mutant ♀’s have abnormal wing postures restricting their escape from rearing cups following eclosion, which reduces survival.

**Video S3. Timelapse of *myo-fem* mutant flight**. Cages consisting of *myo-fem* mutant ♀’s, *myo-fem* mutant ♂’s, WT ♀’s, and WT ♂’s were recorded over 5.5 minutes. The cages were occasionally tapped to stimulate movement.

**Video S4. Timelapse of pgSIT and WT mosquitoes**. Cages consisting of pgSIT mutant ♀’s, *pgSIT*^*♂*^’s, WT ♀’s, and WT ♂’s were recorded for 5.5 minutes. The cages were occasionally tapped to stimulate movement/flight.

**Video S5. DAM assay video**. Short clip of the DAM assay’s monitor tube in action with a WT ♀passing the infrared beam during flight.

**Video S6. Male courtship assay**. pgSIT males are strongly attracted to the female flight tone indicating strong mating behavior.

**Video S7. Model-predicted impact of releases of *pgSIT* eggs in Onetahi, Tetiaroa, French Polynesia**. Time-series for female *Ae. aegypti* population density and elimination probability are depicted for four sample release schemes depicted in **Figure 4**.

## Supplemental Dataset

**File S1. Amplicon EZ sequencing data**.

## References

1. T. C. Winegard, The Mosquito: A Human History of Our Deadliest Predator (Penguin, 2019).

2. C. L. Moyes, J. Vontas, A. J. Martins, L. C. Ng, S. Y. Koou, I. Dusfour, K. Raghavendra, J. Pinto, V. Corbel, J.-P. David, D. Weetman, Contemporary status of insecticide resistance in the major Aedes vectors of arboviruses infecting humans. PLoS Negl. Trop. Dis. 11, e0005625 (2017).

3. D. A. Dame, C. F. Curtis, M. Q. Benedict, A. S. Robinson, B. G. J. Knols, Historical applications of induced sterilisation in field populations of mosquitoes. Malar. J. 8 Suppl 2, S2 (2009).

4. X. Zheng, D. Zhang, Y. Li, C. Yang, Y. Wu, X. Liang, Y. Liang, X. Pan, L. Hu, Q. Sun, X. Wang, Y. Wei, J. Zhu, W. Qian, Z. Yan, A. G. Parker, J. R. L. Gilles, K. Bourtzis, J. Bouyer, M. Tang, B. Zheng, J. Yu, J. Liu, J. Zhuang, Z. Hu, M. Zhang, J.-T. Gong, X.-Y. Hong, Z. Zhang, L. Lin, Q. Liu, Z. Hu, Z. Wu, L. A. Baton, A. A. Hoffmann, Z. Xi, Incompatible and sterile insect techniques combined eliminate mosquitoes. Nature. 572, 56–61 (2019).

5. J. E. Crawford, D. W. Clarke, V. Criswell, M. Desnoyer, D. Cornel, B. Deegan, K. Gong, K. C. Hopkins, P. Howell, J. S. Hyde, J. Livni, C. Behling, R. Benza, W. Chen, K. L. Dobson, C. Eldershaw, D. Greeley, Y. Han, B. Hughes, E. Kakani, J. Karbowski, A. Kitchell, E. Lee, T. Lin, J. Liu, M. Lozano, W. MacDonald, J. W. Mains, M. Metlitz, S. N. Mitchell, D. Moore, J. R. Ohm, K. Parkes, A. Porshnikoff, C. Robuck, M. Sheridan, R. Sobecki, P. Smith, J. Stevenson, J. Sullivan, B. Wasson, A. M. Weakley, M. Wilhelm, J. Won, A. Yasunaga, W. C. Chan, J. Holeman, N. Snoad, L. Upson, T. Zha, S. L. Dobson, F. S. Mulligan, P. Massaro, B. J. White, Efficient production of male Wolbachia-infected Aedes aegypti mosquitoes enables large-scale suppression of wild populations. Nat. Biotechnol. 38, 482–492 (2020).

6. D. O. Carvalho, A. R. McKemey, L. Garziera, R. Lacroix, C. A. Donnelly, L. Alphey, A. Malavasi, M. L. Capurro, Suppression of a Field Population of Aedes aegypti in Brazil by Sustained Release of Transgenic Male Mosquitoes. PLoS Negl. Trop. Dis. 9, e0003864 (2015).

7. K. C. Long, L. Alphey, G. J. Annas, C. S. Bloss, K. J. Campbell, J. Champer, C.-H. Chen, A. Choudhary, G. M. Church, J. P. Collins, K. L. Cooper, J. A. Delborne, O. R. Edwards, C. I. Emerson, K. Esvelt, S. W. Evans, R. M. Friedman, V. M. Gantz, F. Gould, S. Hartley, E. Heitman, J. Hemingway, H. Kanuka, J. Kuzma, J. V. Lavery, Y. Lee, M. Lorenzen, J. E. Lunshof, J. M. Marshall, P. W. Messer, C. Montell, K. A. Oye, M. J. Palmer, P. A. Papathanos, P. N. Paradkar, A. J. Piaggio, J. L. Rasgon, G. Rašić, L. Rudenko, J. R. Saah, M. J. Scott, J. T. Sutton, A. E. Vorsino, O. S. Akbari, Core commitments for field trials of gene drive organisms. Science. 370, 1417–1419 (2020).

8. J. Champer, A. Buchman, O. S. Akbari, Cheating evolution: engineering gene drives to manipulate the fate of wild populations. Nat. Rev. Genet. 17, 146–159 (2016).

9. N. P. Kandul, J. Liu, H. M. Sanchez C, S. L. Wu, J. M. Marshall, O. S. Akbari, Transforming insect population control with precision guided sterile males with demonstration in flies. Nat. Commun. 10, 84 (2019).

10. D. Navarro-Payá, I. Flis, M. A. E. Anderson, P. Hawes, M. Li, O. S. Akbari, S. Basu, L. Alphey, Targeting female flight for genetic control of mosquitoes. PLoS Negl. Trop. Dis. 14, e0008876 (2020).

11. O. S. Akbari, I. Antoshechkin, H. Amrhein, B. Williams, R. Diloreto, J. Sandler, B. A. Hay, The developmental transcriptome of the mosquito Aedes aegypti, an invasive species and major arbovirus vector. G3. 3, 1493–1509 (2013).

12. S. Gamez, I. Antoshechkin, S. C. Mendez-Sanchez, O. S. Akbari, The Developmental Transcriptome of Aedes albopictus, a Major Worldwide Human Disease Vector. G3. 10, 1051–1062 (2020).

13. E. C. Degner, Y. H. Ahmed-Braimah, K. Borziak, M. F. Wolfner, L. C. Harrington, S. Dorus, Proteins, Transcripts, and Genetic Architecture of Seminal Fluid and Sperm in the Mosquito Aedes aegypti. Mol. Cell. Proteomics. 18, S6–S22 (2019).

14. J. Chen, J. Luo, Y. Wang, A. S. Gurav, M. Li, O. S. Akbari, C. and Montell, Suppression of female fertility in Aedes aegypti with a CRISPR-targeted male-sterile mutation. Under Revision at PNAS (2021).

15. S. O’Leary, Z. N. Adelman, CRISPR/Cas9 knockout of female-biased genes AeAct-4 or myo-fem in Ae. aegypti results in a flightless phenotype in female, but not male mosquitoes. PLOS Neglected Tropical Diseases. 14 (2020), p. e0008971.

16. M. Li, T. Yang, N. P. Kandul, M. Bui, S. Gamez, R. Raban, J. Bennett, H. M. SánchezC, G. C. Lanzaro, H. Schmidt, Y. Lee, J. M. Marshall, O. S. Akbari, Development of a confinable gene drive system in the human disease vector. Elife. 9 (2020), doi:10.7554/eLife.51701.

17. M. Li, M. Bui, T. Yang, C. S. Bowman, B. J. White, O. S. Akbari, Germline Cas9 expression yields highly efficient genome engineering in a major worldwide disease vector, Aedes aegypti. Proc. Natl. Acad. Sci. U. S. A. 114, E10540–E10549 (2017).

18. E. C. Degner, L. C. Harrington, Polyandry Depends on Postmating Time Interval in the Dengue Vector Aedes aegypti. Am. J. Trop. Med. Hyg. 94, 780–785 (2016).

19. H. M. C. Sánchez, S. L. Wu, J. B. Bennett, J. M. Marshall, MGDrivE: A modular simulation framework for the spread of gene drives through spatially explicit mosquito populations. Methods Ecol. Evol., 229–239 (2019).

20. E. A. Mordecai, J. M. Cohen, M. V. Evans, P. Gudapati, L. R. Johnson, C. A. Lippi, K. Miazgowicz, C. C. Murdock, J. R. Rohr, S. J. Ryan, V. Savage, M. S. Shocket, A. Stewart Ibarra, M. B. Thomas, D. P. Weikel, Detecting the impact of temperature on transmission of Zika, dengue, and chikungunya using mechanistic models. PLoS Negl. Trop. Dis. 11, e0005568 (2017).

21. J. Bouyer, M. J. B. Vreysen, Concerns about the feasibility of using “precision guided sterile males” to control insects. Nature Communications. 10 (2019),, doi:10.1038/s41467-019-11616-9.

22. H. Briegel, M. Hefti, E. DiMarco, Lipid metabolism during sequential gonotrophic cycles in large and small female Aedes aegypti. J. Insect Physiol. 48, 547–554 (2002).

23. N. P. Kandul, J. Liu, H. M. Sanchez C, S. L. Wu, J. M. Marshall, O. S. Akbari, Reply to “Concerns about the feasibility of using ‘precision guided sterile males’ to control insects.” Nature Communications. 10 (2019),, doi:10.1038/s41467-019-11617-8.

24. B. J. Matthews, O. Dudchenko, S. B. Kingan, S. Koren, I. Antoshechkin, J. E. Crawford, W. J. Glassford, M. Herre, S. N. Redmond, N. H. Rose, G. D. Weedall, Y. Wu, S. S. Batra, C. A. Brito-Sierra, S. D. Buckingham, C. L. Campbell, S. Chan, E. Cox, B. R. Evans, T. Fansiri, I. Filipović, A. Fontaine, A. Gloria-Soria, R. Hall, V. S. Joardar, A. K. Jones, R. G. G. Kay, V. K. Kodali, J. Lee, G. J. Lycett, S. N. Mitchell, J. Muehling, M. R. Murphy, A. D. Omer, F. A. Partridge, P. Peluso, A. P. Aiden, V. Ramasamy, G. Rašić, S. Roy, K. Saavedra-Rodriguez, S. Sharan, A. Sharma, M. L. Smith, J. Turner, A. M. Weakley, Z. Zhao, O. S. Akbari, W. C. Black 4th, H. Cao, A. C. Darby, C. A. Hill, J. S. Johnston, T. D. Murphy, A. S. Raikhel, D. B. Sattelle, I. V. Sharakhov, B. J. White, L. Zhao, E. L. Aiden, R. S. Mann, L. Lambrechts, J. R. Powell, M. V. Sharakhova, Z. Tu, H. M. Robertson, C. S. McBride, A. R. Hastie, J. Korlach, D. E. Neafsey, A. M. Phillippy, L. B. Vosshall, Improved reference genome of Aedes aegypti informs arbovirus vector control. Nature. 563, 501–507 (2018).

25. M. Li, L. Y. C. Au, D. Douglah, A. Chong, B. J. White, P. M. Ferree, O. S. Akbari, Generation of heritable germline mutations in the jewel wasp Nasonia vitripennis using CRISPR/Cas9. Sci. Rep. 7, 901 (2017).

26. D. G. Gibson, L. Young, R.-Y. Chuang, J. C. Venter, C. A. Hutchison 3rd, H. O. Smith, Enzymatic assembly of DNA molecules up to several hundred kilobases. Nat. Methods. 6, 343–345 (2009).

27. H. Li, Minimap2: pairwise alignment for nucleotide sequences. Bioinformatics. 34, 3094– 3100 (2018).

28. M. U. G. Kraemer, M. E. Sinka, K. A. Duda, A. Q. N. Mylne, F. M. Shearer, C. M. Barker, C. G. Moore, R. G. Carvalho, G. E. Coelho, W. Van Bortel, G. Hendrickx, F. Schaffner, I. R. F. Elyazar, H.-J. Teng, O. J. Brady, J. P. Messina, D. M. Pigott, T. W. Scott, D. L. Smith, G. R. W. Wint, N. Golding, S. I. Hay, The global distribution of the arbovirus vectors Aedes aegypti and Ae. albopictus. Elife. 4, e08347 (2015).

